# Cell-to-cell variability in the yeast pheromone response: Cytoplasmic microtubule function stabilizes signal generation and promotes accurate fate choice

**DOI:** 10.1101/093195

**Authors:** C.G. Pesce, S. Zdraljevic, A. Bush, V. Repetto, W. Peria, R.C. Yu, A. Colman-Lerner, R. Brent

## Abstract

In a companion paper, we carried out a high-throughput screen to identify genes that suppressed cell-to-cell variability in signaling in yeast. Two genes affected cytoplasmic microtubules that can connect the nucleus to a signaling site on the membrane. Here, we show that microtubule perturbations that affected polymerization and depolymerization, membrane attachment, and force generation increased variability. For some perturbations, "outlier" cells drove the increased variability. Bypass experiments that activated the PRS ectopically at downstream points indicated that microtubule-dependent processes might stabilize the membrane-recruited scaffold protein Ste5. The variability caused by microtubule perturbations required the MAP kinase Fus3. Microtubule perturbations hindered stable scaffold formation and decreased the accuracy of a polarity-dependent fate choice. Our experiments suggest that membrane-attached microtubules stabilize signaling by scaffold-bound Fus3, and are consistent with a model in which signaling irregularities from changes in microtubule function are amplified by cross-stimulatory feedbacks among PRS proteins. The fact that microtubule perturbations also cause aberrant fate and polarity decisions during embryonic development and cancer initiation suggests that similar variation-reducing processes might also operate in metazoans.

## Introduction

In metazoan tissues, populations of cells must determine extracellular ligand concentrations, and, frequently must determine the ligand gradient vector. Cells then must make appropriate fate decisions (including induction of gene expression, cell cycle arrest, differentiation, and polarity determination) in response to those determinations. Maintaining *coherence* (i.e. limiting cell-to-cell variation) in population responses is critical for the organized sequence of cell and tissue interactions during embryonic development, and for the regulation of cell division and differentiation that defines tissue maintenance in the adult.

In *Saccharomyces cerevisiae*, the apparatus (the pheromone response system, or PRS) that allows the response to one such extracellular ligand, the mating pheromone α-factor, is well understood. Sensing of pheromone concentration and induction of genes at appropriate levels in response to pheromone depends on a set of proteins and events that here we called the *signaling arm* of the PRS (Figure 1a). Determination of gradient vector and growth toward mating partner depends on a second set of proteins and events, here called the *polarity determination arm* of the PRS (Figure 1b). The two arms share protein components localized to the same site on the inner plasma membrane (called, respectively, the signaling site or polarity patch). For example, formation of both the signaling site and the polarity patch depends on the βγ (Ste4-Ste18) subunits of the G protein, which dissociate from the α subunit (Gpa1) after pheromone binds the Ste2 receptor. Moreover, each arm stimulates the other's activity (Figure 1e shows an example of S to P stimulation).

**Figure 1.**
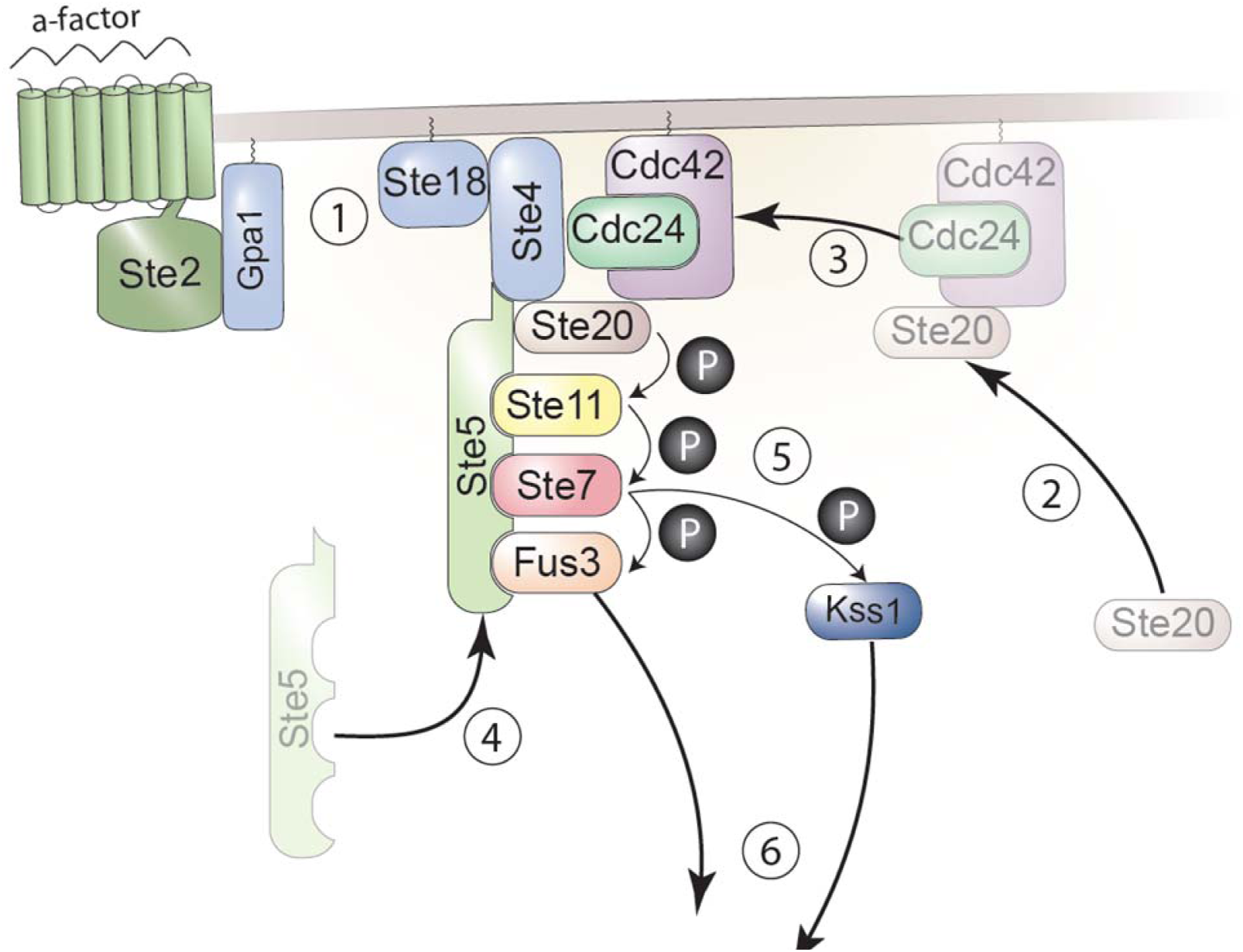
a. The signaling arm of the pheromone response system (PRS). After Ste2 binds the α-factor ligand, the following events lead to the expression of pheromone responsive genes(PRGs). **(1)** The trimeric G-protein Gpa1/Ste4/Ste18 dissociates and thus becomes active. **(2**) Ste20 is recruited to the membrane by a complex containing Cdc42 and Cdc24 (This is an example of positive regulation of the signaling arm by the polarity determination arm, or P to S stimulation) Ste20 makes independent contacts with the membrane (not shown here). **(3)** The Ste20/Cdc24/Cdc42 complex binds to the dissociated G-protein Ste4/Ste18. **(4)** The Ste20/Cdc24/Cdc42/Ste4/Ste18 complex recruits the scaffold protein Ste5 to the cell membrane. **(5)** Thus recruited, Ste5 enables Ste20 activation of the MAP kinase cascade from MAPKKK Ste11 through MAPKK Ste7 and MAPK Fus3. **(6)** Some phosphorylated Fus3 then enters the nucleus to phosphorylate Ste12, Dig1, and Dig2 (not shown), thereby inducing the expression of PRGs, which here include fusions of *P_PRM1_* to fluorescent proteins. Ste7 can also phosphorylate Kss1 which can enter the nucleus and activate PRGs (not shown). Events (1) and (2) are independent and may occur concurrently.

**Figure 1.**
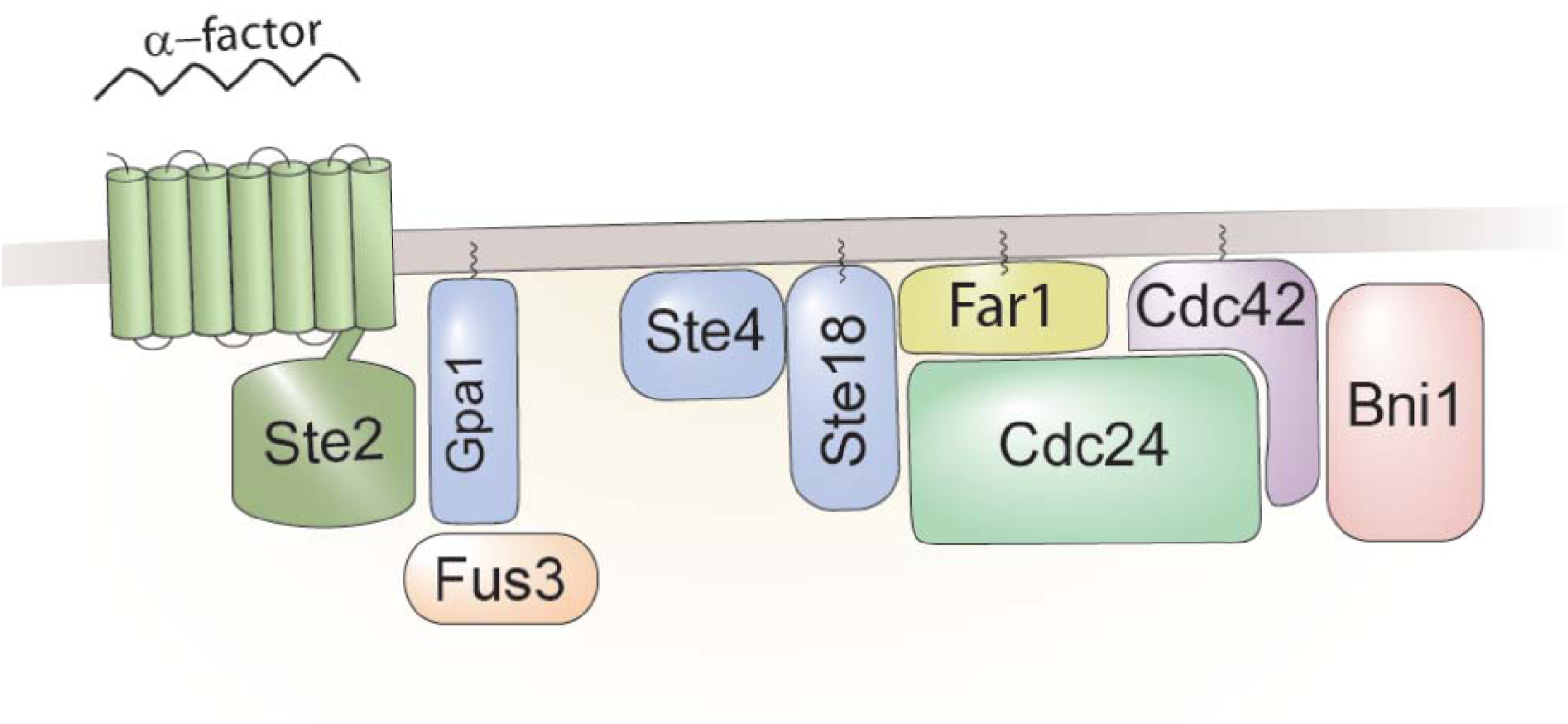
b. Membrane-bound components of the polarity determination arm of the PRS. Many of these proteins (but not Far1 and Bni1) are shared with the signaling arm. Bni1, a formin, stimulates actin cable formation when phosphorylated by Fus3 (Matheos et al. 2004). Actin cables nucleated at Bni1 in turn provides a route for delivery of vesicles and cargo to the membrane and transport and attachment of cytoplasmic microtubules (S to P stimulation, Figure 1e) (Karpova et al. 2000).

**Figure 1.**
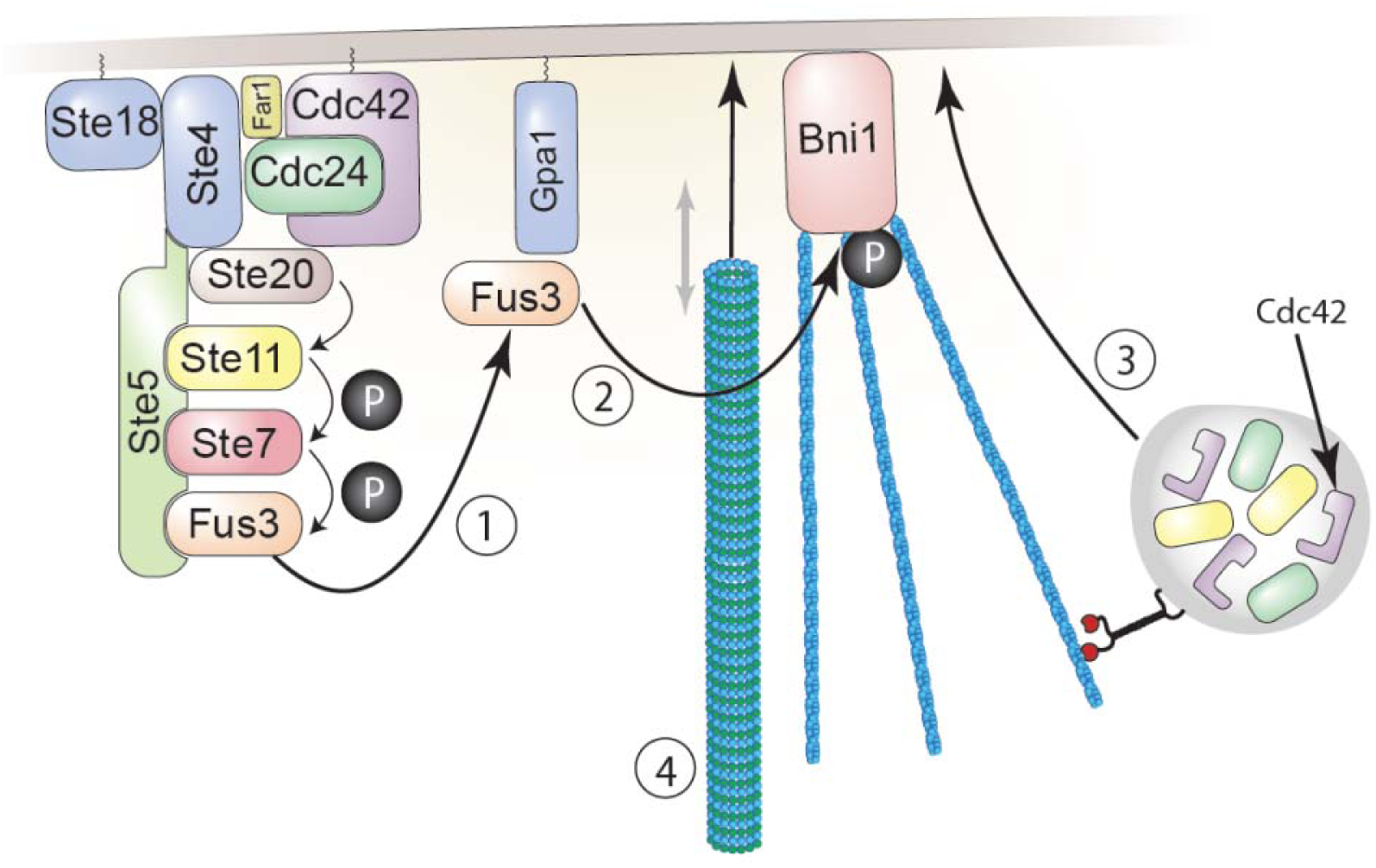
e. Stimulation of polarity patch formation by activity of Fus3 (S to P stimulation). **(1)** Fus3 is phosphorylated and activated due to the operation of the PRS signaling arm. Some phosphorylated Fus3 binds to the Gpa1 subunit of the dissociated G-protein. **(2)** Gpa1-bound Fus3 activates Bni1, which nucleates formation of actin cables. **(3)** Additional proteins involved in signaling, cell polarization, and cell fusion, including Cdc42 and Fus2 (Paterson et al., 2008, not shown) are then trafficked to the membrane as cargo carried along the actin cables. **(4)** Microtubule plus ends are captured by Kar3/Cik1 associated with Gpa1 (not shown) and Kar9/Bim1-complexed plus ends can be walked down the actin cables to the polarity patch by motor proteins including Myo2 (not shown). Microtubules, actin cables and cargo proteins are larger than the approximate scale used here would indicate. Also not shown is a second, more complex instance of stimulation involving Fus3. In this process, activated GPA1 allows the RNA binding protein Scp160 to complex with mRNAs encoding polarity and signaling proteins (called POLs), including Fus3 (Gelin-Licht et al. 2012). These mRNAs are trafficked down actin cables to the site, and, when translated, increase the amounts of proteins such as Fus3 at the site (S to S) stimulation. Activated Fus3 protein stimulates the formation of more actin cables (S to S and S to P stimulation).

Cell-to-cell variation (quantified as η^2^ = (σ/μ)^2^, the square of the ratio between the standard deviation and the mean) in signal, P, transmitted through the PRS signaling arm is under active control (Colman-Lerner et al. 2005, Yu et al. 2008). P is negatively correlated with global gene expression capacity G. The MAPK Kss1 increases η^2^(P) at low pheromone inputs and the MAPK Fus3 decreases η^2^(P) at high inputs (Colman-Lerner et al., 2005). The PRS shows an important "systems level" quantitative behavior, DoRA (<u>Do</u>se-<u>R</u>esponse <u>A</u>lignment) which aligns downstream output with percent receptor occupied (Yu et al. 2008). DoRA can improve the fidelity of information transmission by making downstream responses more distinguishable, and reduce amplification of stochastic noise during signal transmission (Yu et al., 2008, Andrews et al. 2016). The existence of variation suppression mechanisms suggests that, during evolution, accuracy of information transmission by cell signaling systems may have been under selection.

In the companion paper, in order to reveal additional processes that affected variation, we constructed a collection of more than 4,000 yeast strains, each of which carried a deletion of a single nonessential gene. These strains also carried the reporter genes needed to quantify cumulative system output (O), cumulative transmitted signal (P), and variation in these quantities η^2^(O) and η^2^(P)) and output of control promoters in individual cells. We then carried out a large, technically complex, high-throughput (more than 1000 strains with different deletions) flow cytometric and microscope-based screen to find genes that affected cell-to-cell variability in signal transmission and gene induction. From this screen, we identified 50 "variation genes" required for normal cell-to-cell variability in signal and response. Some genes affected variability in system output, η^2^(O), or in transmitted signal η^2^(P), but not the mean in these parameters, O or P. Thus, those two variability phenotypes define independent "axes" of system behavior, and the wild type genes as "canalizing" (Schmalhausen (1949) and Waddington (1942)) the variation (e.g. Jimenez-Gomez et al. 2011, Makumburage and Stapleton 2011) in these traits. Some of these genes that affected η^2^(P) also affected microtubule function. These included *GIM4* and *PAC10/GIM2*, whose products form part of a six-protein prefoldin complex needed for tubulin supply, and *BIM1*, whose product mediates attachment of cytoplasmic microtubule plus-ends to the signaling site. We selected BIM1 and one of the two prefoldin genes, GIM4, as candidates to explore the relationship between microtubule function and signaling variation.

During yeast mating, cytoplasmic microtubules are critical for nuclear fusion (Molk and Bloom, 2006). Early in the pheromone response, Gpa1, the Gα subunit of the heterotrimeric G-protein, associates with the pheromone receptor Ste2, binds the microtubule motor Kar3/Cik1, and recruits it to the signaling site (Zaichick et al. 2009). Kar3/Cik1 subsequently captures plus ends of astral microtubules, which begin growing (by plus end polymerization and depolymerization) from the SPB in the nuclear membrane. The plus ends randomly explore the inner plasma membrane until they are caught by Gpa1. At least during bud formation, other microtubule plus ends reach the signaling site when complexed with Bim1 and its anchoring partner Kar9 (Maddox et al. 2009). When those plus ends are bound by Bim1 and Kar9, Kar9 can contact the motor protein Myo2 (ten Hoopen et al. 2012), suggesting that Myo2 can then walk the Bim1/Kar9-complexed plus ends down the actin cables emanating from the signaling site until they reach the cell membrane. Once at the signaling site, the microtubules alternately bind Bim1 and Kar3/ Cik1. Bim1 acts passively: it only binds to polymerizing microtubules. Therefore, Bim1 bound microtubules become progressively longer. By contrast, when fixed in the membrane, Kar3/ Cik1 hydrolyzes GTP and depolymerizes microtubule plus ends. Membrane bound microtubules therefore alternate between growth and shrinkage (Maddox et al, 2009). Later in the pheromone response, once the cell fusion stage of mating is completed, microtubules from both partners associate across the fusion site and pull both nuclei towards each other, in a process called "nuclear congression", which culminates with the fusion of the two nuclei into a diploid nucleus (Molk et al. 2006, Molk and Bloom, 2006).

Given the above picture of microtubule function during the pheromone response, we considered how BIM1 and prefoldin might be required to diminish variability in the transmitted signals. One idea was inspired by two observations. The first were reports (Maeder et al. 2007, and very recently, Conlon et al. 2016) showing that, in cells exposed to saturating isotropic pheromone, the concentration of active system MAP kinase (phosphorylated Fus3), occurs in a gradient that has its highest value at the shmoo tip, in which the signaling site is located. The other observation was the opposing actions of Kar3/Cik1 and Bim1 described above. The alternation in forces pulling the nucleus towards the shmoo tip (Kar3/Cik1) and pushing it away (Bim1) causes the nucleus to maintain a "fixed" position at the base of the mating protrusion (Maddox et al., 1999, see BOX and Figure 1c). No role had been assigned to this maintenance or homeostasis of nuclear position; the position of the nucleus would not seem to be relevant to the later processes of nuclear congression and fusion. We therefore hypothesized that the precise control of nuclear position within a Fus3 gradient might reduce variability in the strength of the signal transmitted to the nucleus.

**Figure 1.**
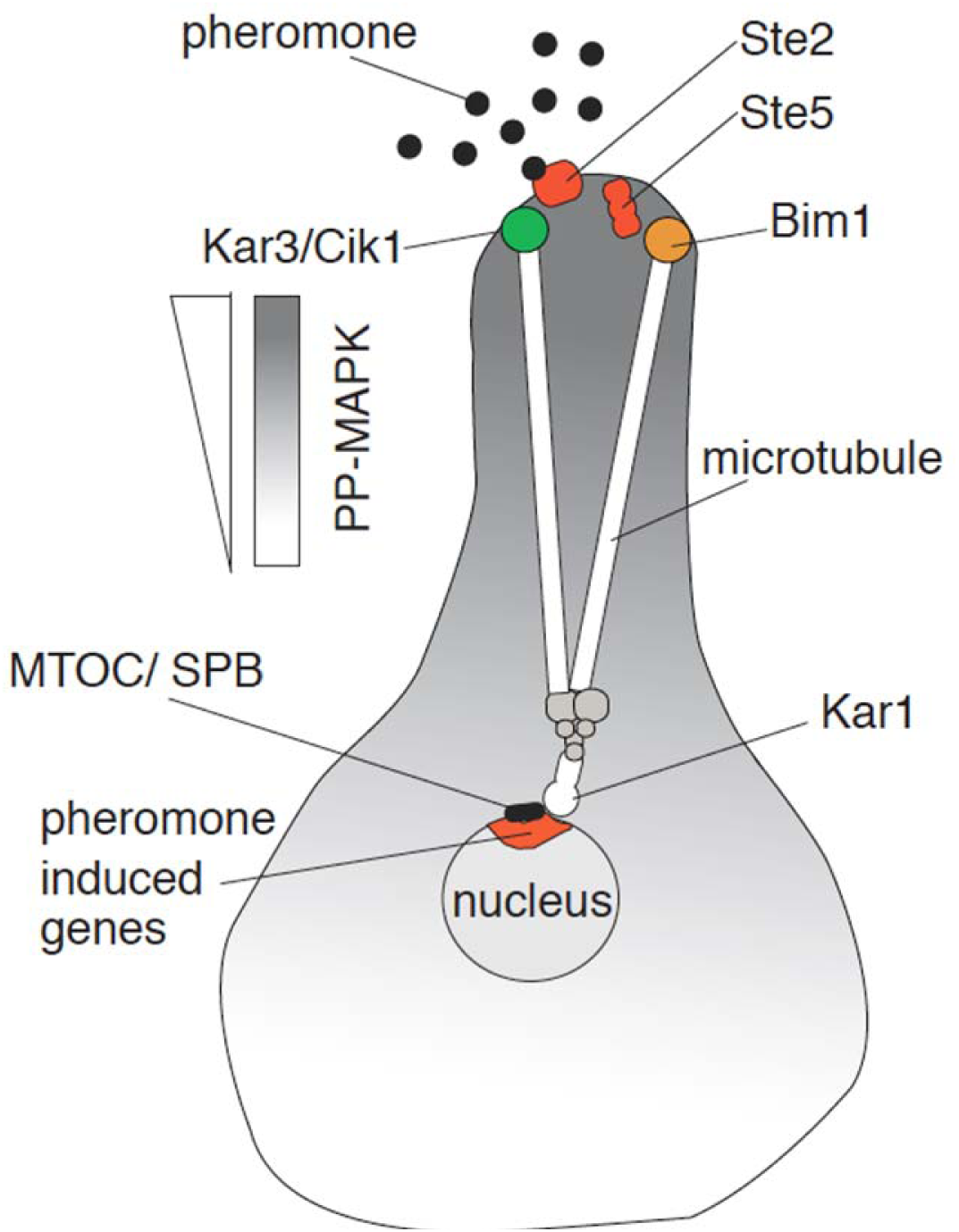
c. Microtubule bridge connecting the signaling site and the nucleus. Schematic of a shmooing cell showing the signaling site (for simplicity, only Ste2 and Ste5 are shown), with Bim1 and Cik1/Kar3 anchors respectively polymerizing (pushing) and depolymerizing (pulling) cytoplasmic microtubules at their plus ends. Cytoplasmic microtubule minus ends attach to the microtubule organizing center/spindle pole body (MTOC/SPB) in a process that requires Kar1. The gradient of grey in the cytoplasm represents a gradient of activated MAPK Fus3 (Maeder et al. (2007), Conlon et al (2016)). Inside the nucleus, the pheromone-induced genes concentrate on the proximal side, closest to the signaling site (Casolari et al., 2005).

A second idea for how normal microtubule function might reduce signaling variability was based on the vulnerability of signaling to positive regulatory interactions at the signaling site/ polarity patch. We wondered if the presence of multiple microtubule plus ends at the signaling site might, perhaps because of their sheer size (the outer diameter of each microtubule is 24 nm) affect delivery of new proteins and mRNAs to the site via actin cables (Botstein et al. 1997), or that the polymerization and depolymerization of plus ends might perturb the operation of some other proteins, for example by affecting the local GTP concentration. We realized that any small fluctuation might be amplified by a complex set of direct and indirect positive feedbacks at this membrane site. Activated Cdc42 exerts direct positive feedback on its own recruitment and activation (Butty et al., 2002, Irazoqui et al., 2003) (Figure 1d). In addition, during polarity determination, Cdc42, whose localization is stimulated by actin cables originating from the polarity patch, also recruits and activates Ste20 ( (Moskow et al., 2000), the PRS MAPKKKK, which in turn leads to the activation of the MAP Kinase Fus3 in the signaling arm (here, called P to S stimulation) (Figure 1a). Moreover, during signaling, the MAPK Fus3, after activation while bound to the Ste5 scaffold, binds Gpa1 and activates the formin Bni1, stimulating actin cable formation and causing transport of additional Cdc42 to the site (S to P stimulation, Figure 1e). Moreover, activated Gpa1 activates an mRNA binding protein Scp160, which transports mRNAs including those encoding Fus3 to the site. Due to these complex cross-stimulatory interactions, each arm of the PRS should exerts positive feedback on its own induction and operation. Positive feedbacks can improve the performance of controlled systems (for example water wheels (Molinié, 1837) servo systems, Hazen, 1934, and electronic amplifiers, Armstrong, 1913, Meissner, 1914, and Franklin, 1913). However, without damping or negative feedback, systems with positive feedbacks are prone to instability (Bennett, 1979). In the PRS, cross stimulation and positive feedback are likely constrained by the fact that both arms compete for components in limited supply. For example, both the S and P arms require the Gβ subunit, Ste4, for their operation, but each cell contains only ~2000 molecules of the Gβ complex even when all ˜7500 molecules of the receptor (Thomson et al., 2011) are bound. We thus hypothesized that in normal cells, the attachment, polymerization and depolymerization of microtubule plus ends might be concerted with other activities at the signaling site, so that self-amplifying fluctuations might be minimized.

**Figure 1.**
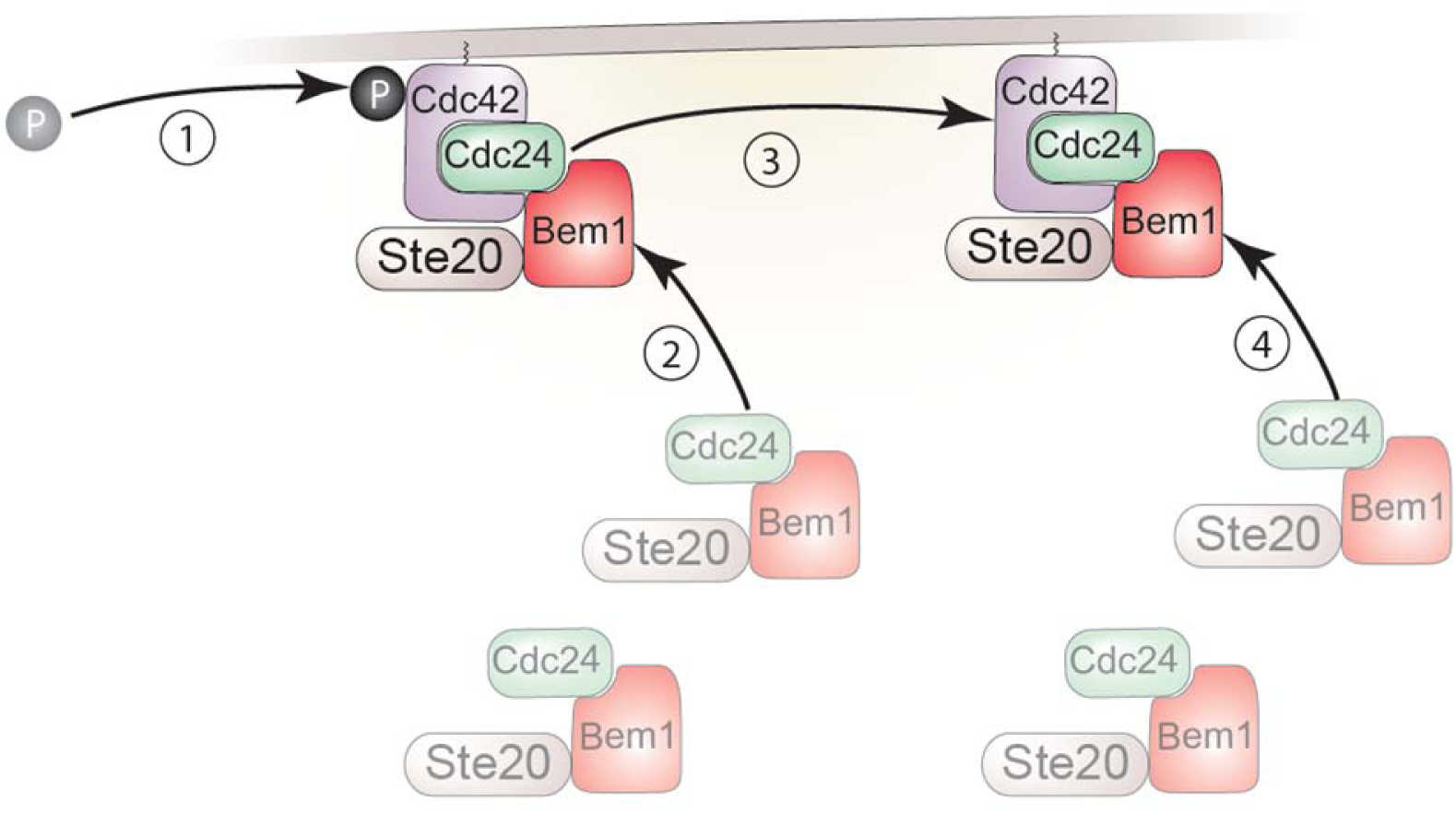
d. Self-stimulation of Cdc42 membrane recruitment and activation. **(1)** The small G protein Cdc42 is activated (converted to GTP-bound form) at some basal stochastic rate. **(2)** Activated Cdc42 recruits a complex containing Bem1, Ste20 and Cdc24. **(3)** Membrane associated Cdc24 (a guanine exchange factor or GEF) activates additional membrane-bound Cdc42. **(4)** As in (2), activated Cdc42 recruits additional Bem1/Cdc24/Cdc42 complexes, completing the positive feedback loop (Kozubkowski et al 2008, Johnson et al. 2011)

Here, we undertook experiments to test these ideas. Our results did not support the idea that nuclear positioning might control signaling variation. Our experiments did show that an attached microtubule bridge reduced variation in signaling and in polarity decisions, that interference with attachment of the bridge impaired the formation and stability of the signaling site, leading to erratic signaling in the PRS. They were thus consistent with the idea that the increased variation in signaling and polarity decisions might be due to instabilities in signaling introduced by microtubule perturbations further amplified by positive feedbacks at the signaling/polarity site.

## Results

### Mutants altered in different variation genes showed distinct differences in signal transmission

From a genetic screen in a companion paper, we found genes that affected cell-to-cell variability in transmitted signal, η^2^(P). Some of these affected microtubule function. These included *GIM4* and *PAC10/GIM2*, whose products are needed for tubulin subunit supply, and *BIM1*, whose product is needed for attachment of plus ends to the signaling site and their subsequent polymerization.

To better characterize signaling in *Δbim1* and *Δgim4* mutants and to test the idea that increased η^2^(P) was due to their failure to position the nucleus correctly within a cytoplasmic gradient of activated Fus3, we constructed strains (SI and Table S1) that carried the fluorescent protein reporter for the pheromone response subsystem (*P_PRM1_* promoter driving mCherry), a constitutive control reporter to allow measurement of activity of the gene expression subsystem, common to all genes (*P_ACT1_* promoter driving CFP), as well as a histone Htb2-YFP chimera to allow precise measurement of nuclear position (see below).

We used time-lapse microscopy in single cells of these strains to measure total system output (O, the accumulated mCherry), transmitted signal (P, the ratio between the accumulated mCherry and CFP), and variability in these values. Figure 2 shows plots of P in single cells over time ("trajectories") after stimulation with 20 nM pheromone. We first analyzed how these trajectories were grouped. The reference strain ("wild-type") cell trajectories were clustered relatively tightly: The most extreme cells deviated from the average P by less than 50%, while in *Δbim1* and *Δgim4* populations, the most extreme cells deviated from the mean by substantially more than 50%. The *Δbim1* and *Δgim4* cells differed strikingly in terms of how common such extremely high or low Ps were. *Δgim4* cell populations presented a broader distribution of trajectories around the mean. In contrast, in *Δbim1* populations, most cells had trajectories like the reference strain, but a number of "outlier" cells showed P very far from the mean.

**Figure 2.**
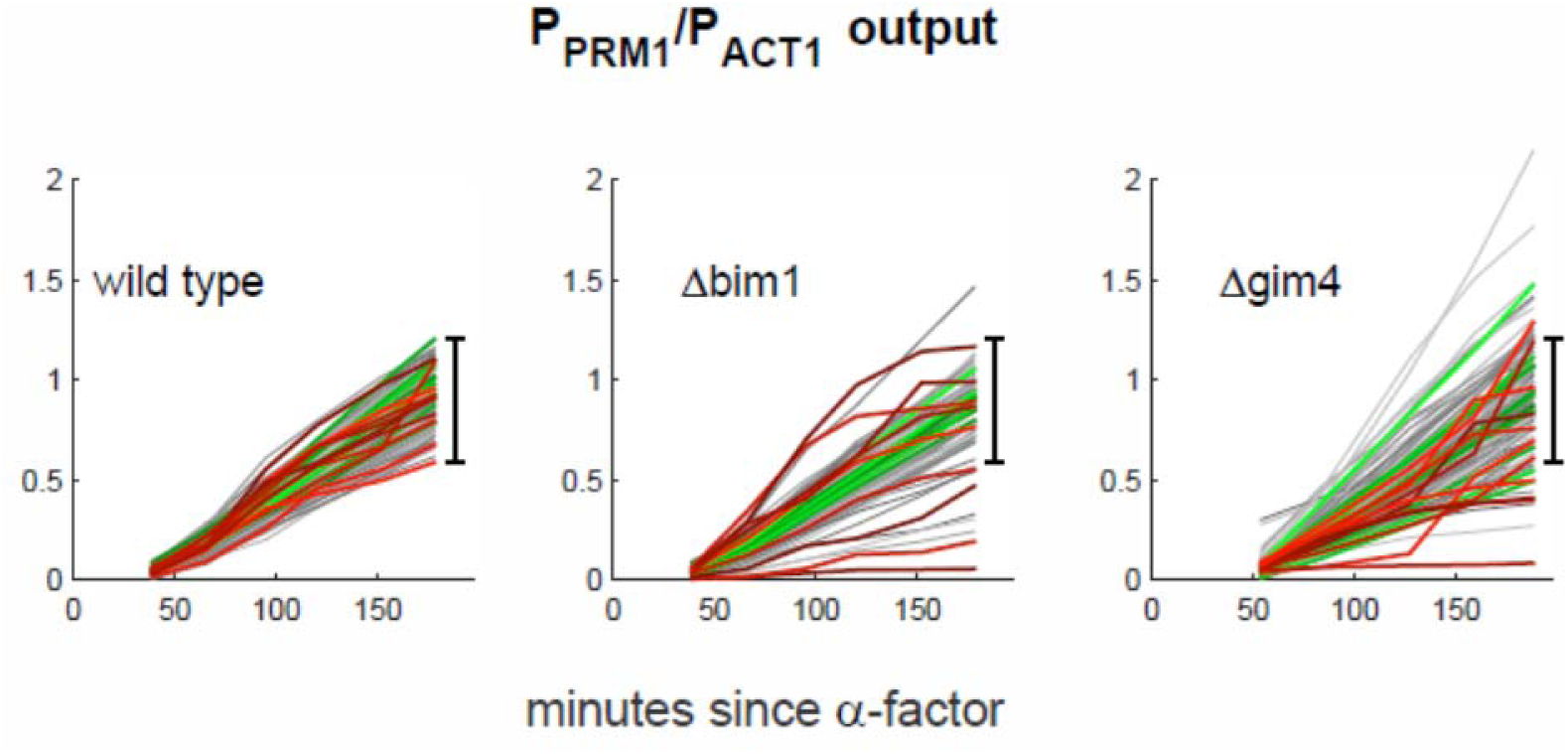
Accumulated signal P vs time in reference and microtubule perturbed cells. y axis shows P measured in individual cells of the reference (GPY4143, "wild type"), *ΔBim1* (GPY4144) and *Δgim4* (GPY4150) strains. We induced the PRS by addition of 20nM pheromone to the medium and imaged cells every 30 minutes as in (Colman-Lerner et al. 2005 and Gordon et al.,2007). In all populations, about 5% of the cells do not respond to pheromone induction. Traces correspond to pathway output (inducible *P_PRM1_-mCherry* signal/ constitutive *P_ACT1_-CFP* signal) from individual cells followed over time. For each strain, variation in the inducible *P_PRM1_-mCherry* signal was larger than variation in the constitutive *P_ACT1_-CFP* signal (not shown). For each strain, we colored the 10 most stable trajectories green, and the 10 least stable (crooked) trajectories red. The black bar shows the full range of WT responses, for comparison.

To quantify these differences and test their significance, the standard deviation or similar statistics are not well suited, since many distinct distributions may have the same standard deviation, and kurtosis is too sensitive to outliers (see Discussion). Thus, we developed a distribution, the progressive spread distribution (PSD), whose median (MPS) is an appropriate measure of the way a distribution is shaped. The MPS value conveys the dispersion observed in the better-behaved half of the cell population. The higher the MPS, the broader the distribution (see Methods). For the cells in Figure 2, the MPS for the reference population was 0.22 (95% CI 0.19 - 0.23), for *Δbim1* cells it was 0.24 (95% CI 0.17 - 0.31, not distinguishable from reference), while for *Δgim4* cells it was 0.36 (95% CI 0.32 - 0.37). Restated, these results show that the distribution of trajectories in *Δgim4* cells is significantly broader overall than WT, while, for *Δbim1* cells, population behavior differs because of outlier cells, while the majority of cells behave normally. We consider the possible significance of these metrics and of outlier cells later (see discussion).

We then analyzed the stability of the trajectories over time. In all populations, the trajectories of most individual cells were stable, while a few cells showed unstable increases in P (evidenced as erratic or "crooked" trajectories (red traces in Figure 2)). For reference cells, stable and crooked trajectories occurred throughout the range of values of P. For *ΔBim1* cells, cells with crooked trajectories occurred at the same frequency above and below the mean P, and accounted for almost all the cells with extreme values of P. By contrast, *Δgim4* cells showed crooked trajectories only for values of P below the mean. Since *Δgim4* cells also showed a reduction in system output (O), this result suggested that the increased variation in P (η^2^(P)) in *Δgim4* populations might be linked to reduced signal, with some cells with low signal exhibiting unstable trajectories. We take crooked trajectories as evidence of erratic operation of the signaling machinery during the course of the experiment (see discussion).

### Microtubule perturbations have distinct effects on the position and movement of the nucleus relative to the signaling site

To pursue the question of whether variation in nuclear position caused variation in transmitted signal, we determined the relation between P and the distance to the nucleus from the signaling site in single cells. In *Δbim1* cells, Bloom and collaborators had measured this distance under saturating pheromone stimulation (Maddox et al. 1999). Based on this and other work (Maddox et al. 2003, Zaichick et al., 2009, Box) we expected that *Δbim1* cells would not be able to properly position their nuclei during pheromone stimulation. However, there was no published work on the effects of the *Δgim4* mutation or other prefoldin mutants on nuclear positioning. Thus, as a next step we sought to learn the effects of microtubule perturbations on nuclear positioning.

We thus measured the position of the nucleus in the same cells in which we analyzed trajectories. To do so, we used the signal from Htb2-YFP, which labeled the nucleus, to measure the shortest distance between the base of the mating protrusion (starting with the site on the cell perimeter at which the protrusion starts forming) and the nuclear edge. We first plotted time courses of nuclear distance from the shmoo base in these cells, stimulated with a saturating dose of pheromone. Figure 3a shows the results. As expected and described (Maddox et al, 1999), in the reference cells nuclei moved synchronously and rapidly from various initial locations to 0-1 μm away from the base of the mating protrusion. In *Δbim1* populations some nuclei moved towards the base of the mating protrusion, while other nuclei seemed to wander in the cytosol. In *Δbim1* cells, both sorts of nuclear movement were characterized by sudden changes in position, more frequent and drastic than in reference cells. In contrast, in *Δgim4* populations nuclear position was as coherent as in reference cells. However, in *Δgim4* cells, nuclei took longer to initiate their movement towards the shmoo base, with the majority remaining at the same distance for two hours or more.

**Figure 3.**
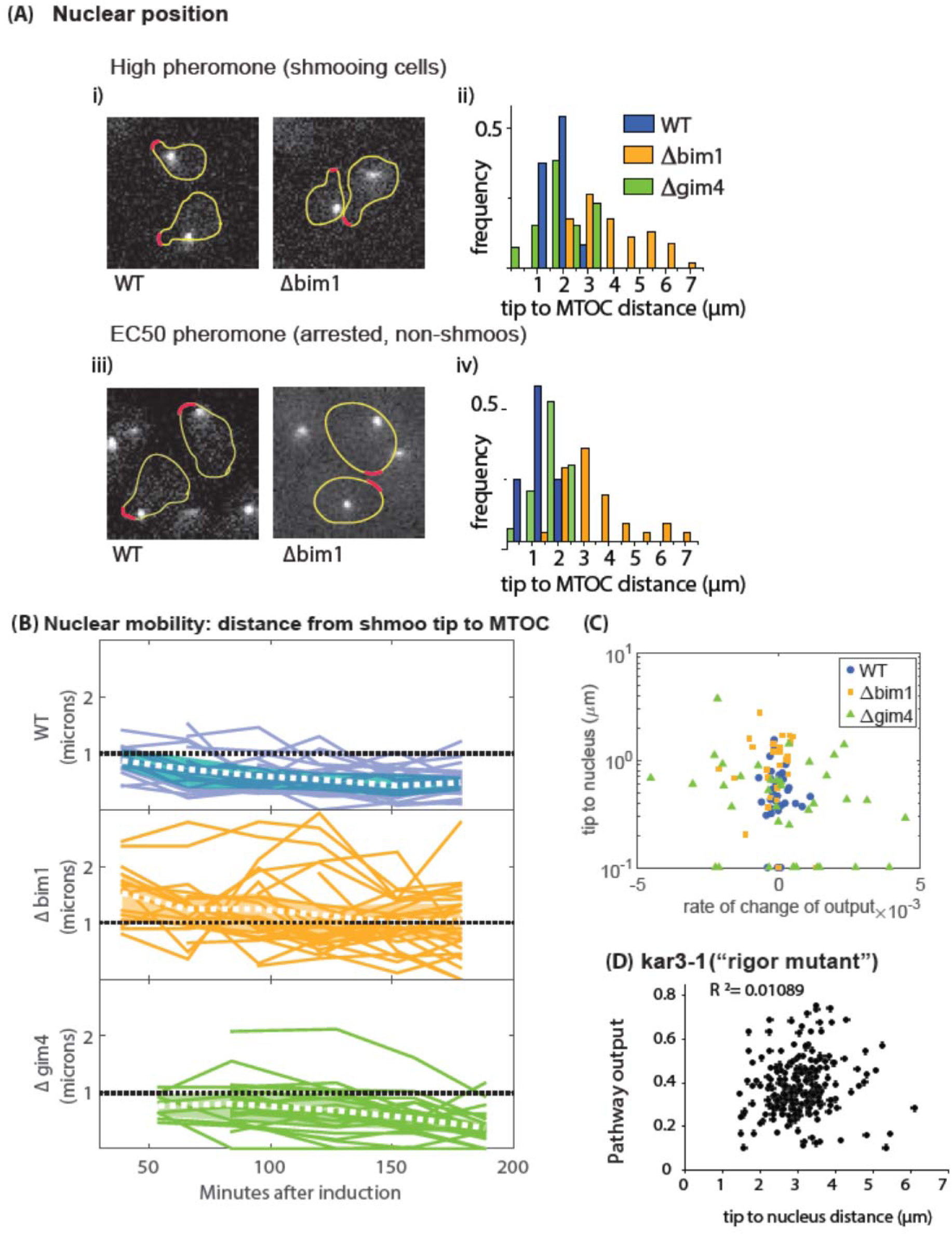
Distance and change in distance between nucleus and signaling site in microtubule perturbed cells. **(A)** Images of cells expressing Spc42-GFP, which marks the MTOC, and Sec8-RFP, which marks the signaling site. On the left, images show reference (“WT”) and *Δbim1* cells (GPY1752 and GPY1709)-- cells of the *Δgim4* strain (GPY1759) were visually indistinguishable from images of WT cells and thus were not shown. Images were recorded after 90 min of exposure to high (20 nM, i-ii) or intermediate (2 nM, iii-iv) pheromone doses. On the right, histograms show distances in μm between the MTOC and the signaling site (the shmoo tip) for reference (WT, blue), *Δbim1* (orange) and *Δgim4* (green) cells **(B)** Time courses of the distance between base of the shmoo (where the cell wall is before the protrusion starts growing) and the nuclear edge (visualized by Htb2-Venus) after stimulation with 20 nM pheromone, for reference, *Δbim1*, and *Δgim4* cells as in A, with the same color-coding. At each time point, we computed the 0.25 and 0.75 quantiles, and used these to shade the middle two quartiles. The white dashed line marks the mean, and we drew a dashed black reference line to indicate 1 μm distance. **(C)** Signaling strength (P) in *kar3-1* (rigor mutant) strains. We stimulated *kar3-1* cells (SGA150, expressing Htb2-Venus to mark the nucleus and bearing *P_PRM1_-mCherry* and *P_ACT1_-CFP* reporters to monitor P) with 20 nM pheromone for 180 min. Panel shows the P vs distance between the site of polarization and the nucleus for *kar3-1* cells, with each dot representing a single individual cell from this clonal population. We estimated Pathway output P from the ratio between the output of the *P_PRM1_* and *P_BMH2_* reporters. At this pheromone concentration, these *kar3-1* populations showed three-fold higher signal variation (not shown).

To better measure the position of the nucleus, we made another set of strains (SI and Table S1). In this set, we expressed an Spc42-GFP fusion, which labels the SPB (Spindle Pole Body, the yeast MTOC, MicroTubule Organizing Center), and a Sec8-mCherry fusion, which labels the signaling site/ polarity patch. We then stimulated these cells with pheromone. The use of these two labels allowed us to obtain precise measures of the distance between the SPB and the signaling site at all pheromone doses (Figure 3aii and iv). However, because the signals were weak and we needed to acquire several images in the z-axis to find them, photobleaching of the fluorophores prevented us from taking time-courses in individual cells. Figure 3ai shows the results. In these strains at saturating pheromone doses, the Spc42-GFP signal appeared as a single dot in the cytosol of each cell, and the Sec8-mCherry signal formed a small crescent at the shmoo tip (Figure 3Ai). At concentrations near the receptor EC_50_, Sec8-mCherry localized to one of the two otherwise indistinguishable ends of the enlarged arrested cells (Figure 3aiii). In reference cells stimulated for two hours at saturating pheromone doses, the nuclei were between 1 to 3 μm from the signaling site (Figure 3aii), in agreement with the work of Maddox et al. (1999). In *Δbim1* cells the mean SPB-to-signaling site was larger, and the distance ranged from between 2 to 7 μm. In *Δgim4* cells, the mean SPB-to-site distance was unchanged relative to the reference cells, but the distribution of distances was broader. In reference cells at the EC_50_ pheromone dose, which causes cell cycle arrest but not shmooing (Figure 3A iii), we observed an even shorter distance between the nucleus and the signaling site, 0 (too close to resolve) - 2 μm, including a striking ~30% of cells in which we could not measure any distance between the nucleus and signaling site (Figure 3A iv). At the EC_50_ dose, effects of the *Δgim4* and *Δbim1* mutations on nuclear distance were qualitatively similar to those at saturating pheromone, with the exception that the *Δgim4* cells showed larger SPB to nuclei distances, in the 1-3 μm range.

Taken together, these results were consistent with the idea that microtubules comprise a force-generating tether connecting the signaling site and the nucleus. They also demonstrated that the *Δbim1* and *Δgim4* mutations partially disrupt its function. They confirmed previous observations about Bim1's role in positioning the nucleus during the pheromone response and showed that prefoldin activity is also required for this process. Thus, both mutations we had identified as causing increases in η^2^(P) also caused changes in one or more aspects of the position of the nucleus relative to the signaling site, including average distance, variation in distance, and tempo of positioning.

### Microtubule-perturbation-induced changes in nuclear position do not correlate with changes in pathway output

We next sought to test the hypothesis that the amount of transmitted signal depended on nuclear position. If so, P in single cells might be a negative (or positive) function of the nucleus-to-signaling-site distances. We examined this correlation in populations of reference and mutant cells. Figure 3c shows the results. Neither mutant (*Δgim4* or *Δbim1*) nor reference cells show a correlation between instantaneous or average nucleus to site distance and the rate of change of output O (here, a proxy for P). The most straightforward interpretation of these results is that the distance between the signaling site and nucleus does not affect the strength of the transmitted signal, P.

These results did not exclude the idea that the lack of relation between P and the nucleus-signaling site distance might be because the nucleus is constantly moving, especially in the mutant strains, and as a result of that movement, the observed P is actually an average of higher and lower Ps (when the nucleus is closer or farther from the signaling site, respectively). To test this possibility, we generated a *kar3-1* (rigor mutation) Htb2-YFP strain in which the microtubule bridge was stable and did not change in length. The Kar3-1 protein binds microtubules plus-ends, but can neither actively depolymerize nor release them. *kar3-1* strains therefore maintain cytoplasmic microtubules attached both the shmoo tip and nucleus, and these microtubules do not change in length (Maddox et al., 2003; Meluh and Rose, 1990). In our kar3-1 strain, the nucleus did not move to the shmoo base, its distance from the signaling site varied significantly from cell to cell (1.5 to 6 μ, Figure 3d), but it did not change over time in each cell. Figure 3D shows that, in *kar3-1* cells, there was no correlation between nucleus to signaling site distance and output O, a proxy for P. This result confirmed that P is not a function of the distance between the nucleus and the signaling site, and showed that the MAPK signal can reach the nucleus even when the nucleus is not in line with the signaling site.

### Inhibition of microtubule polymerization and depolymerization increased signaling variation

The above results showed that increased signaling variation in *Δbim1* and *Δgim4* strains were not related to the position of the nucleus relative to the signaling site, and were consistent with the idea that their effects were due to alteration of some other aspect(s) of microtubule function. To explore how altered microtubule function might increase variation, we examined the effect of perturbations that disrupted different aspects of microtubule function.

We first tested chemical treatments that cause the depolymerization of dynamic microtubules. We used the microtubule-polymerization inhibitors benomyl and nocadazole at concentrations reported to induce breakdown of cytoplasmic microtubules (Palframm et al. 2006). Figure 4 (lower left) shows that chemical disruption of microtubules caused a decrease in P at doses of pheromone higher than the EC50, consistent with the decrease in P in *Δbim1* and *Δgim4* strains. However, unlike the effects of *Δbim1* and *Δgim4*, these microtubule poisons did not affect η^2^(P) (Figure 4, upper left). This result was again consistent with the idea that the effect of *Δbim1* and *Δgim4* on η^2^(P) could be due to these mutants perturbing specific aspects of microtubule function. In this view, microtubules are present and functional in both deletion strains, but, are either impaired in their ability to attach to the site of polarized growth (*Δbim1*) or less abundant (*Δgim4*). However, these results again do not exclude the idea that the effects of *Δbim1* and *Δgim4* on η^2^(P) could be unrelated to their effects on microtubule function. To distinguish between these possibilities, we used additional genetic perturbations that affected other aspects of microtubule function.

**Figure 4.**
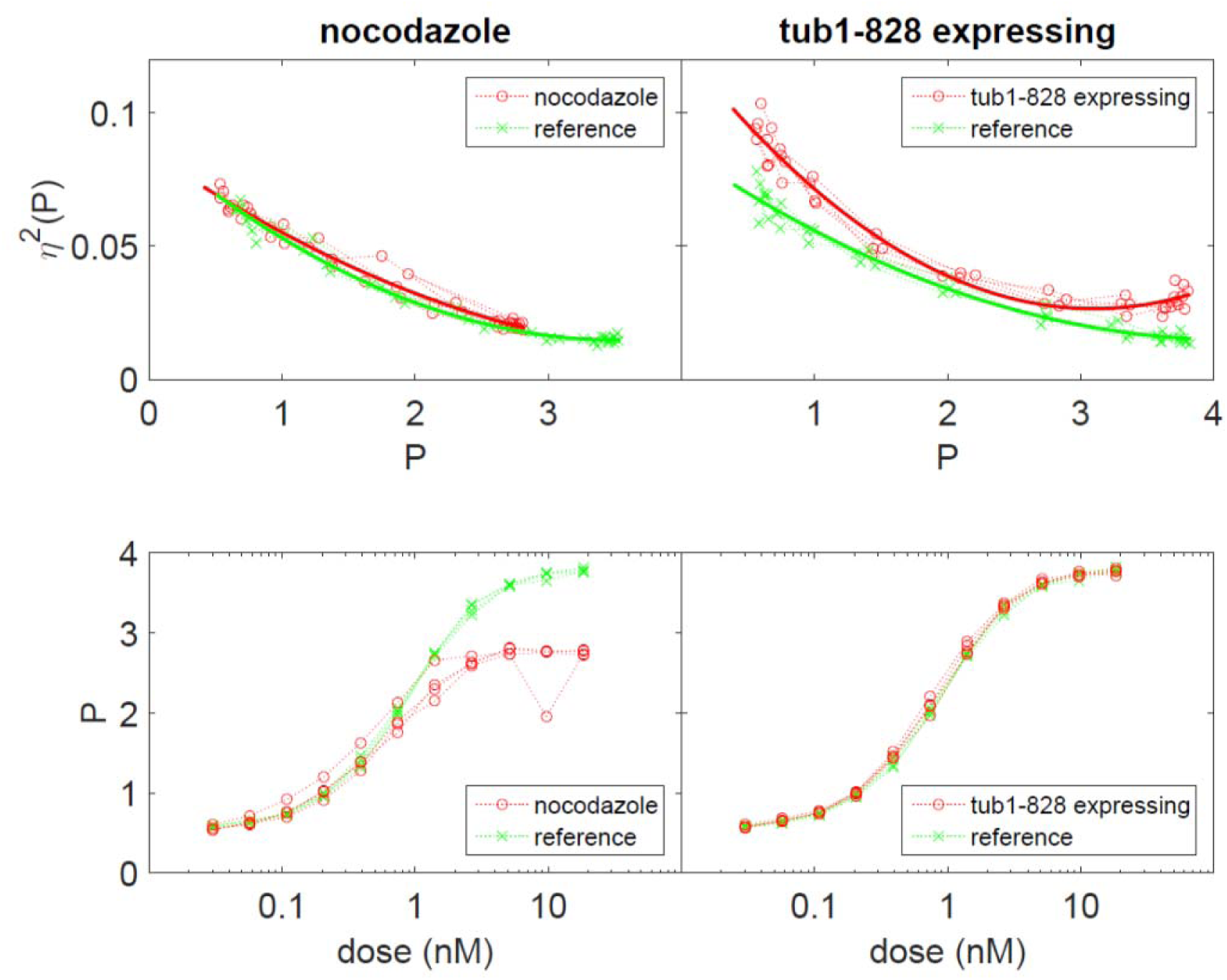
Cell-to-cell variability in transmitted signal, η^2^(P), and in cumulative transmitted signal, P, in strains treated with chemical and genetic inhibitors of microtubule polymerization and depolymerization. Top row shows η^2^(P) as a function of P (top row), Bottom row shows P as a function of dose. The *Tub1-828-expressing* strain (GPY1873) was a derivative of GPY4000. Strains in left column were treated with nocadazole/ benomyl, strains in right column were and perturbed by *Tub1-828 expression* (right column).

### Microtubule perturbations affecting membrane attachment, polymerization and depolymerization increased signaling variation

We next disrupted microtubule polymerization and depolymerization by a targeted genetic perturbation: expression of a dominant negative variant of α-tubulin, encoded by the *TUB1-828* allele. When expressed at low levels in cells containing wild type α-tubulin, Tub1-828 is incorporated into the growing microtubule tip, where it blocks polymerization and depolymerization (Anders and Botstein, 2001). Since high levels of Tub1-828 block further cell division (Anders and Botstein, 2001), we used the *GAL1* promoter, expressed at low levels in cells grown in glucose medium (Guarente et al., 1982, Yocum et al. 1984), to drive Tub1-828 expression (SI and Table S1). Figure 4 shows that, in glucose medium, Tub1-828 expression increased variation in transmitted signal, η^2^(P) at all pheromone doses tested. In galactose medium, at higher levels of Tub1-828 expression where cells were blocked from further division, we also observed increased η^2^(P) (not shown). We also observed increased η^2^(P) in cells in which higher levels of Tub1-828 were expressed in an inducible system in which estradiol activates *P_GAL1_* (see below and (Louvion et al. 1993)). These results strongly supported the idea that altering microtubule function increases variation in transmitted signal.

We next blocked microtubule pulling by constructing (SI and Table S1) otherwise-isogenic *Δkar3* and *Δcik1* strains and measured the effect of these genetic alterations on signal. Kar3 and Cik1 proteins are needed for active, ATP-dependent depolymerization of membrane-attached microtubule plus ends. Figure 5 shows the results. Compared to the reference strain, cells of the *Δkar3* strain showed the same level of output O, while *Δcik1* strains showed a reduction in O, i.e., the measured data cover a smaller horizontal range for *Δcik1* than for the reference strain. For comparable values of O, both perturbations significantly increased variation in transmitted signal, η^2^(P). These results show that two independent mutations that, like *Δbim1*, reduce the ability of microtubules to bind to the site of polarized growth, also increase η^2^(P). Because both *Δbim1* and *Δkar3* also substantially reduced the fraction of cells with a microtubule bridge attached to the signaling site (Maddox et al., Bloom 1999), we hypothesized that the reason *BIM1+* and *KAR3+* cells maintain low variability might be that in these cells the microtubule bridge between nucleus and the signaling site was intact. In this view, the additional microtubule functions of pulling in (*KAR3*) and pushing out (*BIM1*) the nucleus might not be not required for the control of η^2^(P). To test this hypothesis, we again made otherwise-isogenic strains that carried *kar3-1*, the rigor variant of Kar3 in which the microtubule bridge is intact but in which microtubules cannot change in length. Figure 5 shows the results. The rigor mutation slightly lowered O, but increased η^2^(P) by at least a factor of three, as much as the full *Δkar3* lesion. These results show that attachment of the microtubule bridge was not sufficient to control η^2^(P). They suggest that, to lower η^2^(P), microtubules not only have to be attached, but must also be able to alternatively push and pull on the nucleus and exert force.

**Figure 5.**
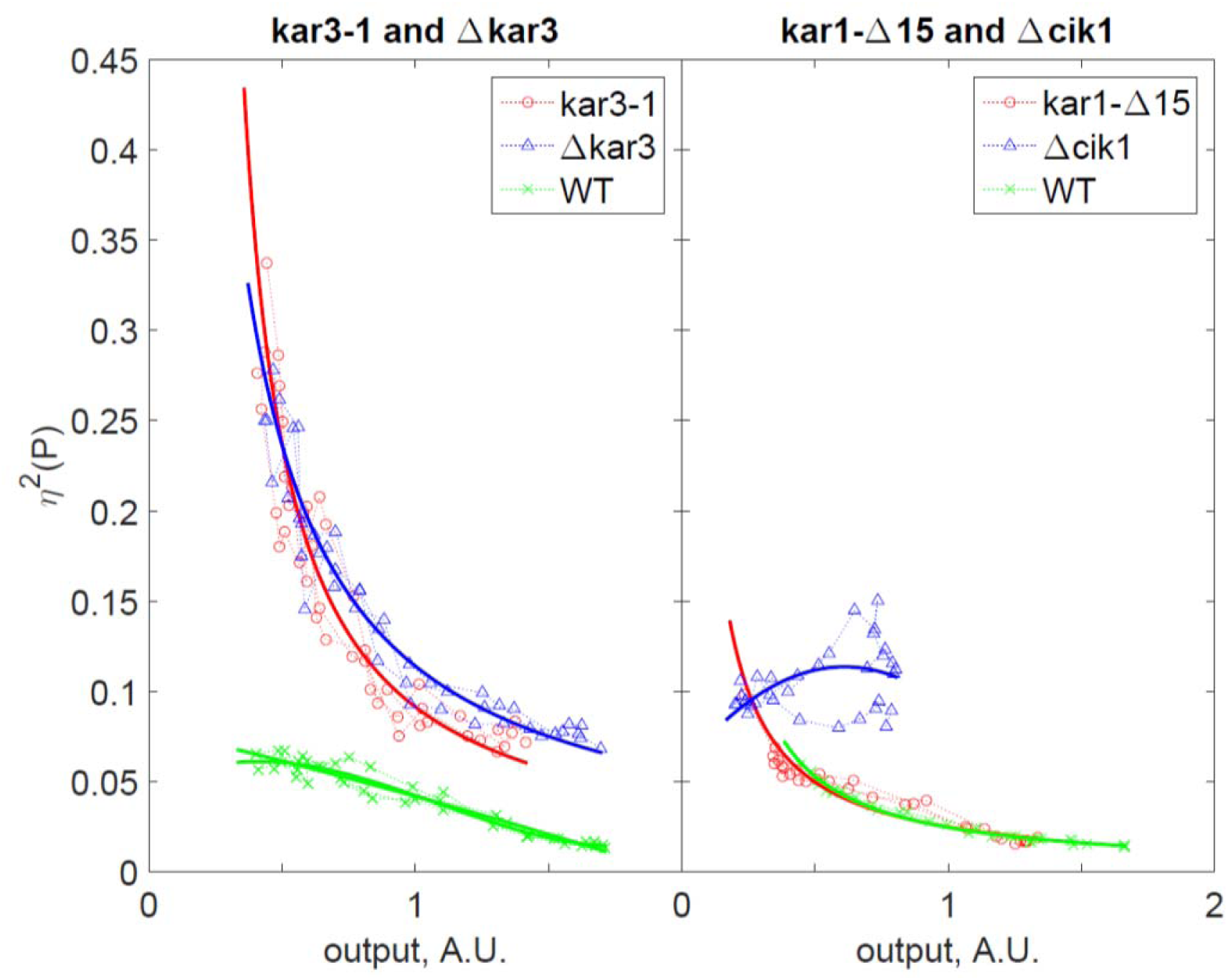
Cell-to-cell variability in signal transmission in cells with mutations affecting microtubule end function. Left panel sows shows η^2^(P) versus output O in *Δkar3* (SGA2015), *kar3-1* (SGA108, the "rigor mutant"), and reference cells (SGA103). Right panel shows η^2^(P), versus O in *kar3-Δ15* (SGA109) cells, *Δcik1* (GPY4123) cells,, and reference cells (SGA103).

Finally, we tested the effects of preventing the attachment of microtubules to the SPB. To do that, we replaced *KAR1* with the *kar1-Δ15* allele (SI and Table S1). *KAR1* encodes a component of the SPB. The *kar1-Δ15* allele expresses a C-terminally truncated variant of Kar1. In pheromone treated cells, the Kar1 C terminus is the main site of minus-end microtubule anchoring to the MTOC (Pereira et al., 1999), and its absence leads to detachment of cytoplasmic microtubules from the nucleus and from the signaling site (Erlehman et al. 2012). Relative to the reference cells, populations of *kar1-Δ15* cells showed decreased O but no change in η^2^(P) (Figure 5), similar to the effect of microtubule depolymerizing drugs (Figure 4). These results support the idea that complete loss of the microtubule bridge reduces P but does not increase η^2^(P).

Taken together, these results suggested that a) an intact microtubule bridge was needed for the full signal transmission (P) by the PRS; b) normal cell-to-cell variability in PRS signaling required correct operation at the plus ends (perhaps the ability to generate force), where the microtubules attach to the signaling site, but not at the minus ends, where they attach to the nucleus. Thus, one or more steps that take place at the signaling site might be affected when microtubules operation is not normal. In the next section, we sought to find the affected steps.

### Genetic bypass of Ste5 recruitment restored coherent population responses

To better understand possible mechanisms by which normal operation of cytoplasmic microtubules might control variation in the pheromone response, we used artificial "bypass" activators to identify pathway step(s) at which microtubule action affected signaling variation. Bypass activators activate systems "ectopically", that is at different steps along the causal chain of signal transmission in the PRS. (Beadle and Ephrussi, 1936,1937, Botstein and Maurer, 1982, Cairns et al. 1992, Takahashi and Pryciak 2008). In a linear pathway, a result showing that a bypass activator is not affected by a perturbation is considered evidence that a perturbation such as a mutation acts upstream or at the ectopically activated step. In systems with feedback and feedforward loops such experiments by themselves cannot preclude other interpretations (Avery and Wasserman, 1992), for example that a downstream perturbation might transmit via feedback its effects upstream of the ectopic activator.

To perform the bypass analysis, we generated a set of strains that conditionally expressed, in the absence of pheromone, one of four different proteins that activated the PRS at different steps, (Takahashi and Pryciak, 2008, SI and Table S1). Expression of each PRS activator protein was driven by *P_GAL1_*, whose activity was controlled in turn by an estrogen-responsive Gal4 derivative, originally developed by Picard and collaborators (Picard 1994). In the order of their function in the pathway, the activator proteins were a) native Ste4, whose expression mimics dissociated (i.e., active) Gβγ dimers (Pryciak and Huntress, 1998); b) Ste5-CTM, a fusion of the Ste5 scaffold with a transmembrane domain, whose expression mimics membrane-recruited (i.e. activated) Ste5 (Pryciak and Huntress, 1998); c) a Ste11-4-Ste7 chimera, a fusion of the Ste11-4 mutant protein, a constitutively active form of Ste11, to the MAPKK Ste7, whose expression mimics activated Ste7 (Harris et al., 2001); and finally d) native Ste12, whose overexpression mimics activated Ste12 (perhaps due to titration of inhibitory factors (Dolan and Fields, 1990; Takahashi and Pryciak, 2008; Tedford et al., 1997)). To prevent interference from basal activation of the native PRS, strains expressing activators b), c) and d) also carried a *Δste5* mutation. Finally, strains in this set also carried both PRS responsive (*P_PRM1_-mCherry*) and constitutive (*P_BMH2_-YFP*) reporters. From each bypass activator strain, we generated four strains which were either unperturbed, carried the *Δbim1 or Δgim4*, or expressed the Tub1-828 protein.

Figure 6A shows the results. Each plot shows η^2^(P) as a function of P at different estradiol doses for the unperturbed strain (gray curves) and the strains with three microtubule perturbations (blue curves). Each row of plots corresponds to one of the four bypass activators (Ste4, Ste5-CTM, Ste11-4-Ste7 and Ste12), while each column of plots corresponds to one of the microtubule perturbations (*Δbim1, Δgim4*, and *Tub1-828-expression*). If the site(s) of action for the microtubule perturbations were downstream of the Ste4 activation step, then the perturbations would still show their effects in the Ste4 activator strains. This was so. All three perturbations *Δbim1, Δgim4* and *Tub1-828 expression* caused increases in η^2^(P), the same effects they had when we stimulated the PRS with pheromone (data not shown). The increase in η^2^(P) in *Δbim1* was similar in both systems, while for *Δgim4* it was less pronounced in the artificial activator system. For *Tub1-828 expression* the increase in η^2^(P) was also similar in both systems, although we could only assess it at low values of P, because *Tub1-828* expression had an unexpectedly strong inhibitory effect on P (see below).

**Figure 6.**
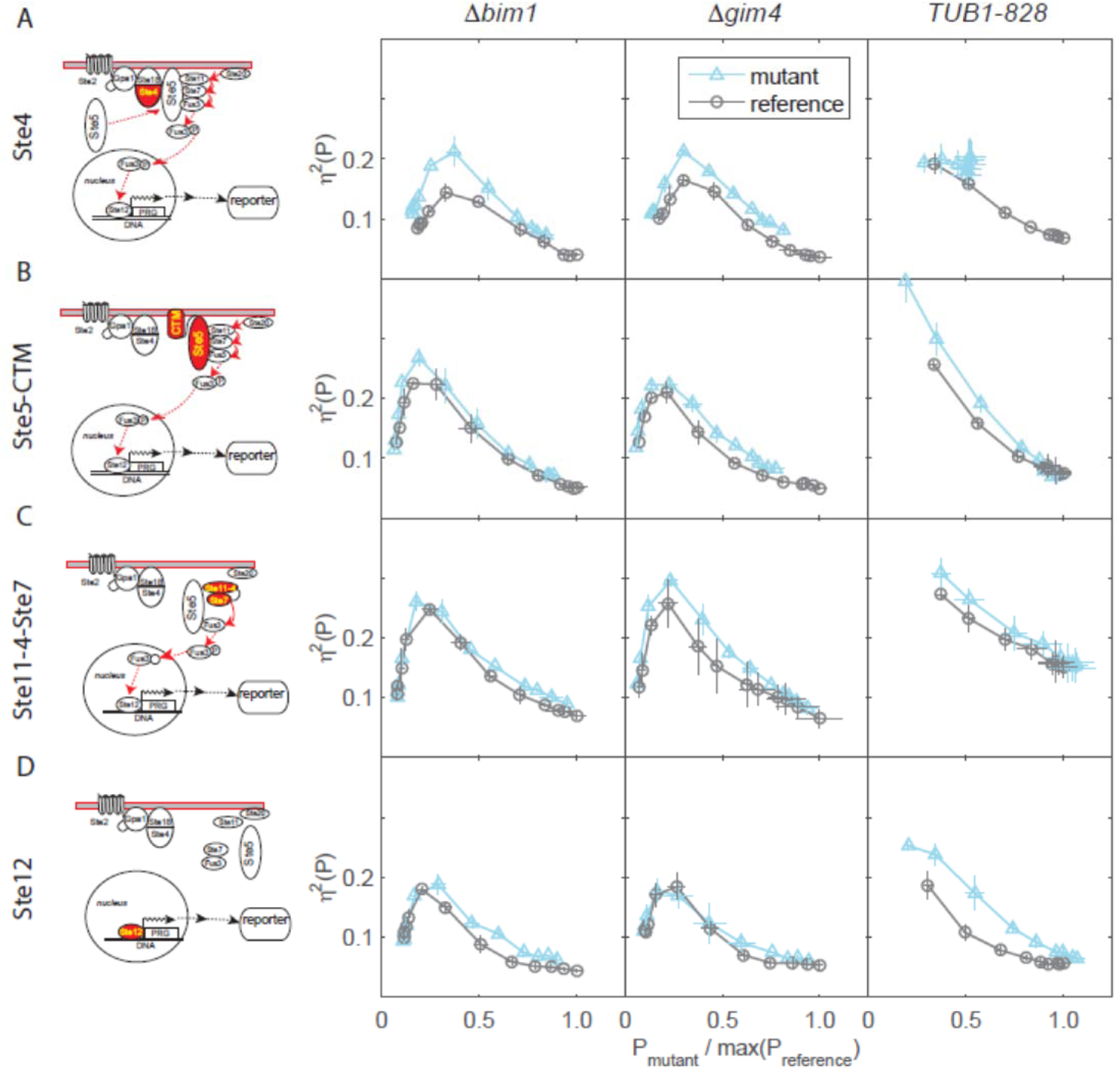
Microtubule perturbations affect cell-to-cell variation in cell signaling at the Ste5 recruitment and MAP kinase steps. We exposed reference ("WT"), *Tub1-828 expressing*, *Δbim1* and *Δgim4* derivatives of GPY1810 (bearing the chimeric genes *P_PRM1_-mCherry*, *P_BMH2_-YFP*, and a gene constitutively expressing the chimeric transcription factor *P_BMH2_-GAL4BD-hER-VP16)*, to the indicated concentrations of estradiol to induce expression of the ectopic activators of the pheromone response system Ste4 (A), Ste5-CTM (B), Ste11-4-Ste7 (C) and Ste12 (D) for 180 min. Cartoons show the proteins of the PRS signaling arm (most shown in Figure 1a) with the ectopic activator highlighted in red. We quantified η^2^(P) from measurements of the two reporter signals in flow cytometry as described. y axis shows cell-to-cell variation in P, uncorrected for η^2^(γ) (η^2^(P)+η^2^(γ)), and x axis shows signal strength P, normalized by the maximum P observed for each reference strain, for cells in three replicate cultures of each mutant and reference strain.

In the *Δbim1* and *Δgim4* strains, the higher-than-reference η^2^(P) phenotype was suppressed by all activators downstream of Ste4 (compare plots in Figure 6A columns 1 - *Δbim1*- and 2 -*Δgim4*)). The suppression was clearest for *Δbim1*, for which the increase in η^2^(P) was present in the Ste4 activator strain and completely suppressed in all downstream activators. The suppression was less conclusive for *Δgim4* because the increase in η^2^(P) in the Ste4 activator strain was smaller and the suppression by the downstream activators was not complete. By contrast, the increase in η^2^(P) caused by *Tub1-828* expression was not suppressed by any of the bypass activators. These results suggested that the microtubule-dependent process(es) perturbed by the *Δbim1* mutation, and possibly also by *Δgim4*, increased signaling variation the PRS at or upstream of the recruitment of Ste5 to the membrane.

We also sought to map the steps in the PRS at which perturbations to microtubule function affected total transmitted signal, P. Figure 6B plots P vs dose. *Δbim1, Δgim4* and *Tub1-828 expression* caused decreases in P. For *Δbim1* and *Δgim4*, the effects on P in the artificial activator system were qualitatively similar to their effects on P in the normal PRS. (see companion paper). The magnitude of the decrease in P due to *Δbim1* in the artificial activator system was similar to that in the PRS. For *Δgim4* the decrease in P in the artificial system was less pronounced. By contrast, in the artificial system, *Tub1-828 expression* a strong inhibitory effect on P, which it did not have in the PRS (Figure 4 and data not shown), perhaps due to very high expression of the Tub1-828 protein at high estradiol doses driven by the Gal4-ER-VP16 fusion.

**Figure 6.**
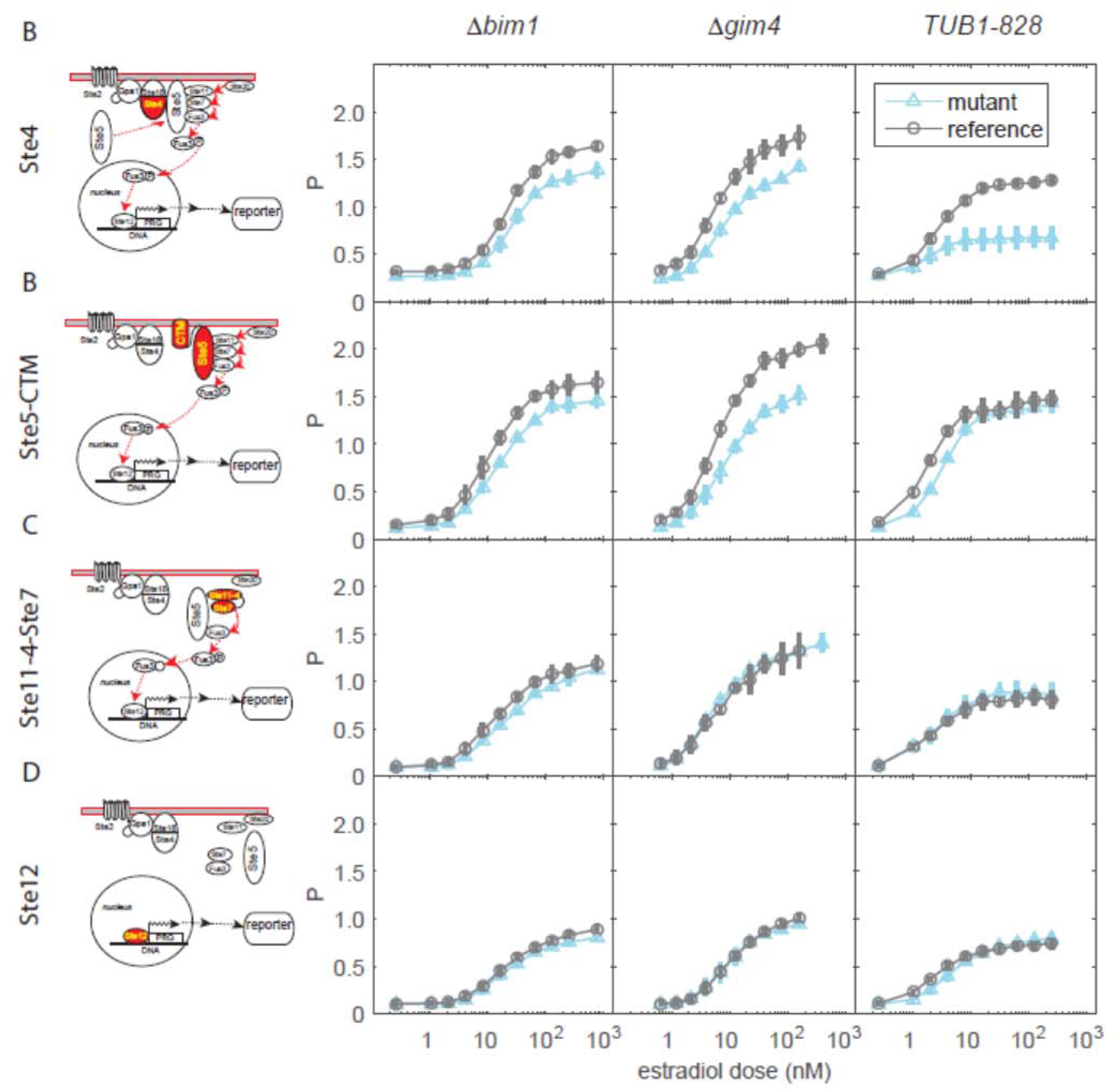
Microtubule perturbations affect transmitted signal P. These are the x-axis values of Figure 6A, plotted here vs. pheromone dose. Reductions in these values are thus also reflected in ranges of P which are reduced relative to reference, for data from mutant strains in the plots of Figure 6A.

The reduction in P caused by *Tub1-828 expression* was completely suppressed by the artificial recruitment of Ste5 in the STE5-CTM strain. That is, the reduction in P caused by *Tub1-828 expression* mapped to the Ste5 recruitment step. By contrast, the reduction in P caused by the *Δbim1* and *Δgim4* mutations was present in the Ste5-CTM strain but bypassed in the strains with downstream activators.

These results showed that, with one exception (very high levels of *Tub1-828 expression)*, the effects of all the perturbations on η^2^(P) and P can be bypassed. They thus established that the microtubule perturbations directly affected the pheromone response by interfering with specific steps in signaling rather than acting indirectly by causing unrelated changes in the cell. They also suggested that microtubule perturbations affected output P and pathway variation η^2^(P) by two different mechanisms, one at or upstream of Ste5 recruitment and the other upstream of the MAPK activation step, and that BIM1 and GIM4 are involved in both of them.

### Induced signaling variation by microtubule perturbations requires Fus3

The above bypass experiments suggested that both the function of the microtubule bridge in normal pathway output and the involvement of Bim1 and Gim4 in maintaining low variability depended on Ste5 recruitment. Ste5 is a cytosolic protein that, upon pheromone stimulation, is recruited to the membrane, primarily by binding to activated Ste4 and secondarily by binding to phospholipids in the membrane. At the membrane, Ste5 localizes the MAPK Ste11 MAPK and the MAPKK Ste7. We showed in previous work that a negative feedback requiring the downstream MAPK Fus3 diminishes retention of Ste5 at the membrane (Yu et al. 2008, Bush and Colman-Lerner 2013). Here we tested whether low variability and normal pathway output mediated by Bim1, Gim4, and the microtubule bridge depended on the action of membrane-localized Ste5 and the partially redundant Fus3 and Kss1 MAPK kinases.

To do so, we generated four strains (SI and Table S1) with a normal PRS, but each bearing deletions of *GIM4* or *BIM1* and also deletions of *FUS3* or *KSS1*, and measured η^2^(P) as a function of P. Figure 7 shows the results. Notably, in *Δbim1* and *Δgim4* strains, the *Δfus3* deletion suppressed completely the increased η^2^(P) observed in the single deletion strains, at all doses (Figure 7). That is, Fus3 protein was required for these microtubule perturbations to increase η^2^(P) (Figure 7).

**Figure 7.**
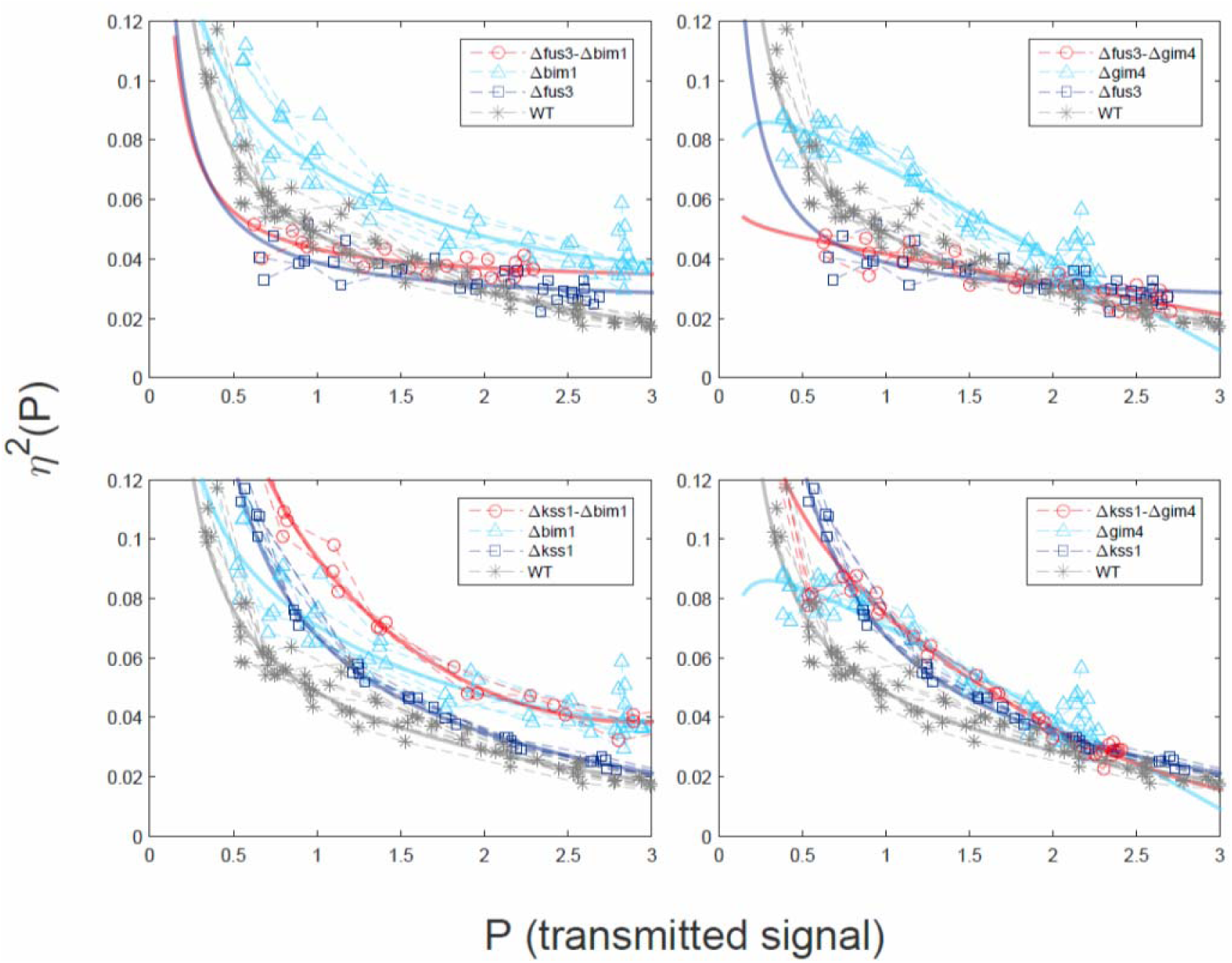
Fus3 is required for microtubule perturbations to increase signal variation. y axis in each panel shows η^2^(P) as a function of P. **Top left.** *Δfus3 Δbim1, Δbim1, Δfus3*, and reference strain (wt). **Top right.** *Δfus3 Δgim4, Δgim4, Δfus3*, and reference (wt). **Bottom left.** *Δkss1 Δbim1, Δbim1, Δkss1*, and reference (wt). **Bottom right.** *Δkss1 Δgim4, Δgim4, Δkss1* and reference strain (wt). *Δbim1* and *Δgim4* strains show increased η^2^(P) relative to WT (p-values are both less than 10^-5^). *Δbim1 Δfus3* and *Δgim4 Δfus3* strains, show WT η^2^(P) (p-values are 0.341 and 0.095, respectively). However, *Δbim1 Δkss1* and *Δgim4 Δkss1* cells show higher η^2^(P) relative to WT (p-values are both less than 10^-5^). The double mutant *Δbim1 Δkss1* had significantly higher η^2^(P) than either *Δkss1* or *Δbim1* alone (p-values are both 10^-5^ or less), while the double mutant *Δgim4 Δkss1* had essentially the same η^2^(P) as either *Δkss1* or *Δgim4* alone (p-values are 0.341 and 0.106, respectively). This shows that Fus3 protein is required for increased η^2^(P).

The fact that the *Δfus3* deletion eliminated the increases in signaling variation caused by the *Δgim4* and *Δbim1* mutations again shows that the increased variation is not a secondary consequence of a generalized increase in variability in cells with disrupted microtubule function, but rather reflects an effect of these mutations on operation of the PRS.

### Perturbation of microtubule function increases variation in establishment and maintenance of the initial signaling site

The above experiments indicated that Ste5 membrane recruitment/retention and the action of membrane bound Fus3 was the point at which *Δbim1* and *Δgim4* deletions introduced variation into signaling. We thus sought to validate these results by directly observing Ste5 membrane localization in the natural PRS and in the perturbed strains. We constructed otherwise-isogenic strains that expressed Ste5 fused in frame to three copies of YFP (Ste5-YFP-YFP-YFP, here called Ste5-YFP). In previous work, we used this construct to show that, in cells in the G1 phase of the cell cycle, cytosolic Ste5-YFP translocates to the plasma membrane isotropically within 2-3 minutes of exposure to isotropic pheromone, followed after 2 to 30 minutes by clustering of the Ste5-YFP signal into the signaling site, a single patch in the plasma membrane that coincides with the site of future shmoo growth (Figure 8A) (Ventura et al.,2014). In addition to the reference strain, this strain set was also comprised of strains that had the *Δbim1*, *Δgim4* or the *Tub1-828 expression* perturbations. We exposed these cells to a high pheromone dose and monitored them by fluorescence microscopy for up to three hours.

**Figure 8.**
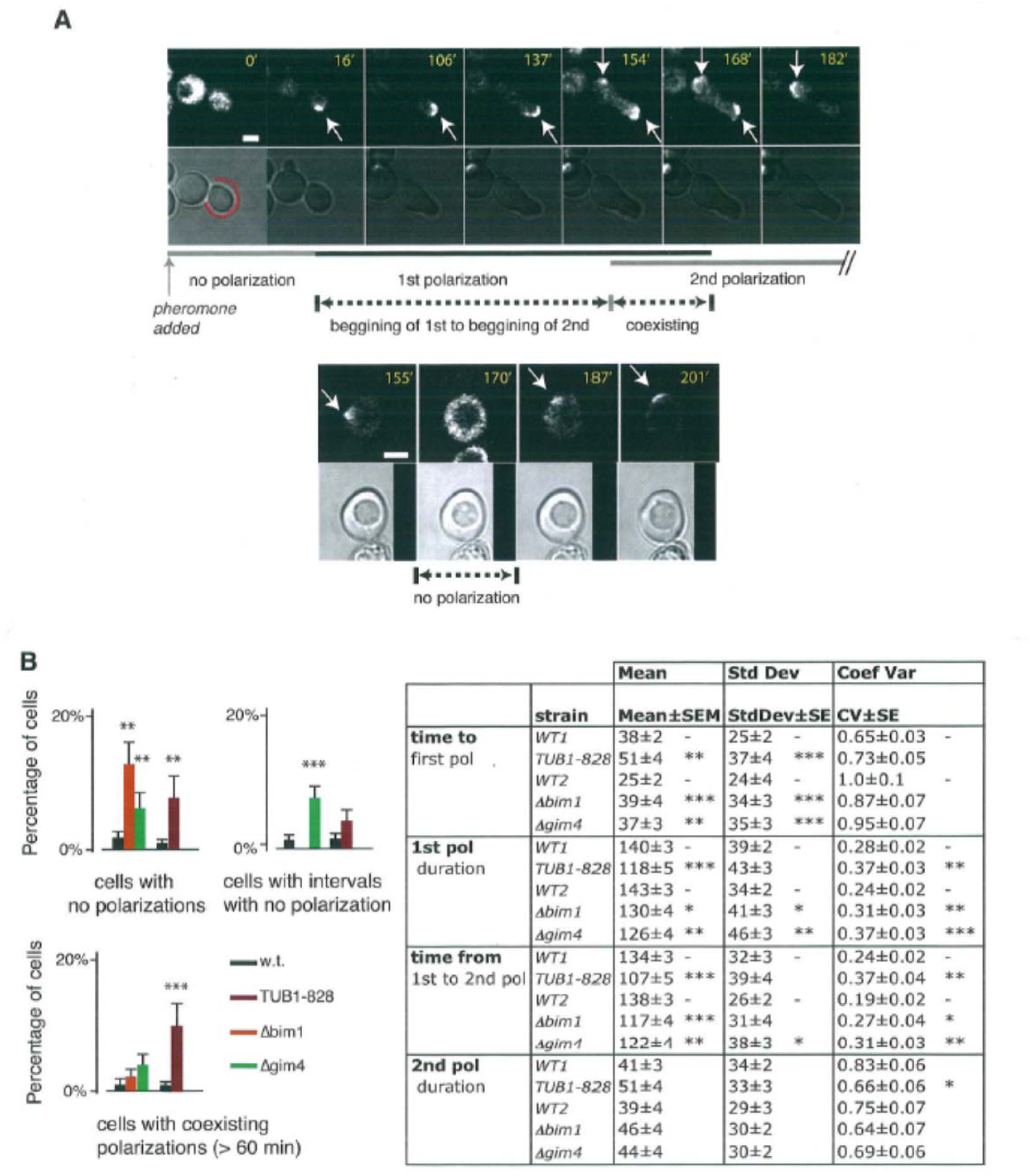
Microtubule perturbations cause Ste5 patches to form less reliably, delay patch formation, and cause patches to persist for less time. We stimulated reference ("WT"), *Tub1-828-expressing*, *Δbim1* and *Δgim4* derivatives of MW003 (bearing 3 copies of *P_STE5_-STE5-3xYFP*, Ventura et al. 2014) with 1μm pheromone and imaged them over time for up to 3.5 hours. **A.** Example of a cell with two co-existing Ste5 patches (top) and a cell with an interval without detectable Ste5 patches between the first and second patch (bottom), marking the dynamic features we quantified: time to 1^st^ patch (polarization), duration of 1^st^ and 2^nd^ polarization, interval in which 1^st^ and 2^nd^ polarization overlap, or gap between them, and time from 1^st^ to 2^nd^ polarization. **B.** Bar graph plots show qualitative defects observed: cells with no polarization, cells with gaps between 1^st^ and 2^nd^ polarizations and cells with overlapping (coexisting) polarizations. Table shows quantitative defects. Data corresponds to the mean +/- SEM, the standard deviation +/- SE and the coefficient of variation (mean divided by the standard deviation) +/- SE. Values for the probability p of the observed data under the null hypothesis that each mutant strain is no different from WT are shown by asterisks: * = p<0.05; ** = p<0.01; *** = p<0.001.

In cells of the reference strain, for the first 45-120 min, the Ste5-YFP patch was visible as a crescent at the tip of the growing shmoo. In most cells the shmoo tip ceased growing after 45-120 min. In these cells a second site of polarized growth appeared later in a different location. Before the second site of growth was morphologically visible, a second Ste5-YFP patch appeared at the site that would became the second site of growth. Progressively, the second site became brighter while the first site faded (Figure 8A). Compared with cells of the reference strain, cells bearing microtubule perturbations showed several common differences. First, in the perturbed strains, a greater number of cells failed to form a signaling site (Figure 8B). This result is in agreement with our earlier observation that in *Δbim1* and *Δgim4* populations about 5% of the cells did not induce a *P_PRM1_-mCherry* reporter after pheromone treatment (Pesce et al., companion paper). Second, in perturbed cells that formed a signaling site, the mean time to site formation was longer and more variable. Third, in perturbed cells, the first site on average lasted a shorter and more variable time. Fourth, in perturbed cells, the time from initiation of the first patch until appearance of the second was on average shorter and more variable. By contrast, compared with the reference strain, the perturbed strains showed no differences in the average and variability of duration of the second patch. Restated, these observations showed that cells with microtubule perturbations had difficulties establishing and maintaining their first signaling sites, but not their second. Some cells failed to form sites, others showed delays of variable length in formation of the first site, and all showed shorter times (also of variable length) to its dissolution. The fact that formation and maintenance of the second site in these cells was normal shows that these three microtubule perturbations do not cause a generalized cellular malaise that indirectly impacts signaling site formation, but that rather that they affected the formation of the first signaling site. The results also suggest that the formation and stability of the second site do not require the microtubule functions we assessed above, and thus suggesting that it did not require the microtubule bridge.

### Perturbation of microtubule function decreases accuracy of polarity decisions in speed mating experiments

Our results pointed to a role for the microtubule bridge in promoting the rapid establishment and sustained maintenance of the signaling site. Haploid *S. cerevisiae* prefer to mate with partners expressing wild type levels of mating pheromone than with partners expressing lower levels (Jackson and Hartwell, 1990). In mixed species mating experiments, haploid *S. cerevisiae* prefer to mate with one another, but also out compete cells from other related species (Murphy and Zeyl, 2010, Murphy and Zeyl, 2012), in part by mating more quickly (Murphy et al.2006). For *cerevisiae*, one part of mating more quickly would be rapid establishment of a stable signaling site oriented toward the correct, high expressing suitor. To test the importance of initial site establishment in mating partner choice, we modified a technique first developed by Hartwell and collaborators (Jackson et al. 1990). In both the original and our methods, mating occurs in mixtures in which *MATa* cells from the strain being tested are combined with equal numbers of two strains of *MATα* partners that i) secrete different amounts of pheromone and ii) carry a marker than can be measured in the resulting diploids. In the variant we used here, one of the *MATα* partner strains (*MATα Δade2 MFALPHA1*) secreted normal levels of pheromone while the other (*MATα ADE2+ Δmfalpha1*) lacked one of the two genes encoding pheromone and thus produced an estimated 5% of WT levels of pheromone. Mating of *MATa Δade2* cells with low-producing suitors produces *Δade2/ADE2+* diploids, which form white colonies on low-adenine medium, while mating of *MATa Δade2* partners with the high-producing suitors produces *Δade2/Δade2* diploids, which form red colonies on the same medium. We mated *MATa* cells with a 10-fold excess of a 1:1 mix of high-producing and low-producing *MATα* partners for different lengths of time, resuspended cells in starvation medium, sonicated the mating mixtures, spread the cells onto selection plates, and scored the color of the resulting diploid colonies.

Figure 9 shows the outcome. In these assays the reference *MATa* cells showed significant preference for mating with the high-pheromone producing partner (65% of the diploid colonies were red, 35% were white.) Among the mutant strains, only *Δgim4* mated with the same discrimination efficiency as wild-type, with *Δbim1*, *Δbim1 Δgim4*, and *Tub1-828-expressing* strains all showing at least marginally significant large reductions in frequency of choosing the normal-pheromone partner, relative to wild-type. We made this comparison via the two-sample t-test; p-values were 0.064, 0.67, 0.025, and 0.034, for the *Δbim1*, *Δgim4*, *Δbim1Δgim4*, and *Tub1-828-expressing* strains, respectively, for the data shown in Figure 9B.)

**Figure 9.**
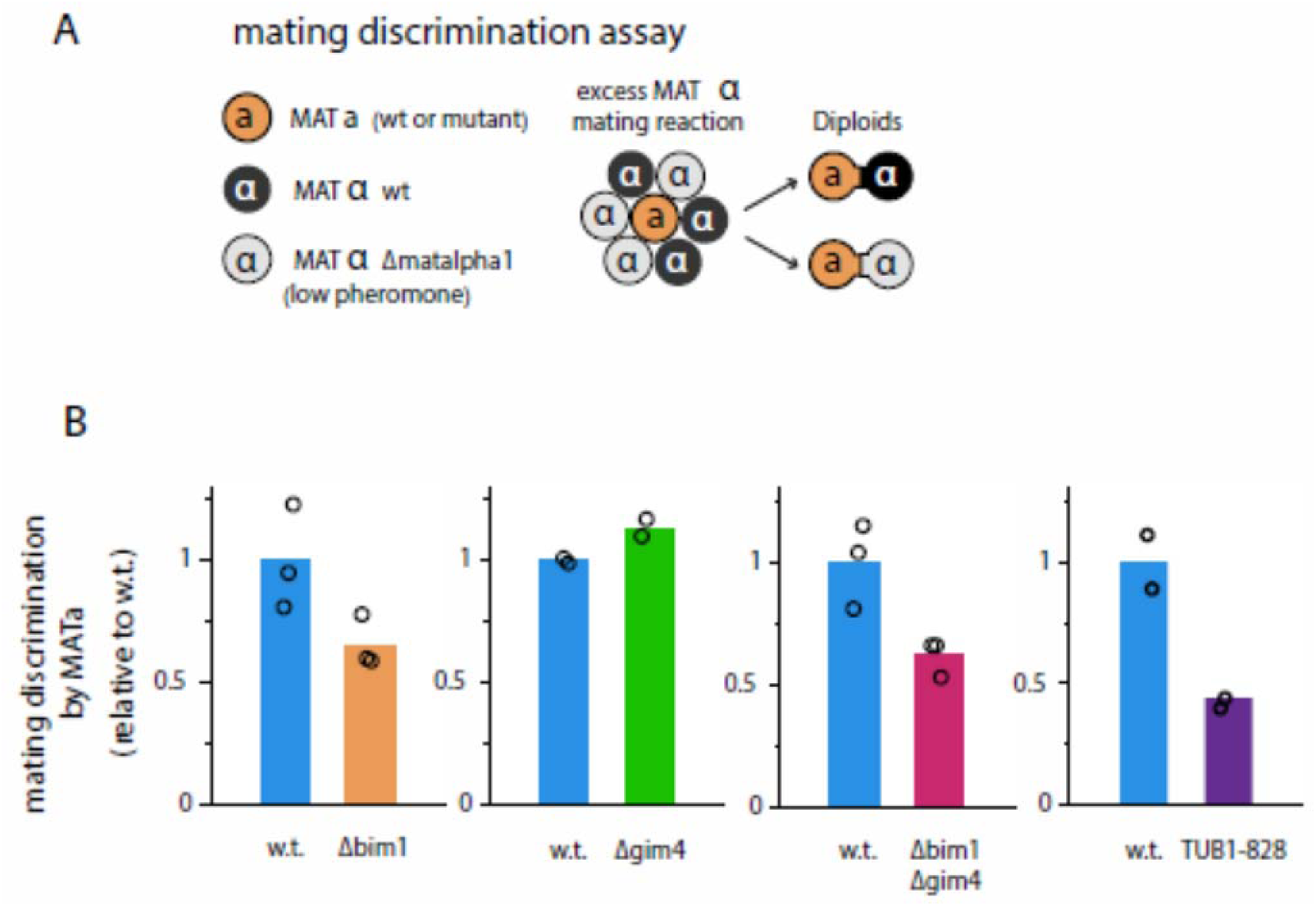
Disrupting microtubule plus end function impairs mating partner choice. **A.** Schematic of discrimination assay, in which MATa cells choose between potential MATα mating partners producing normal levels of α-factor and potential *Δmatalpha1* partners that expressed about 5% as much α-factor. **B.** We performed rapid mating discrimination assays as described in Methods for reference (GPY4078, *Δade2*, and also expressing P_ACT1_-YFP, here called w.t), and for *Δbim1, Δgim4, Δbim1 Δgim4*, and *Tub1-828-expressing* GPY4078 derivatives. We allowed mating to proceed for 5 minutes and then halted it by transferring the cells to medium without nitrogen. Graph shows the discrimination efficiency for three biological replicates of each strain (circles) and the corresponding average (bars). For each strain, we calculated this value as (1-((*Δmatalpha1*/total) diploids / (*Δmatalpha1*/total) haploids)) expressed relative to the value for GPY4078. Loss of Bim1 protein and *Tub1-828-expression* increase variability in transmitted signal η^2^(P), and impair mating partner choice, while loss of Gim4 only increases variation in η^2^(P). In all cases, P is P_PRM1_-XFP/ P_ACT1_-YFP.

In the control matings, done for 1 hour (not shown), the *Δbim1, Δgim4* and *Tub1-828-expressing* strains were as able as the reference strain to mate efficiently and preferentially with high expressing partners, confirming that these perturbations did not disrupt those aspects of mating. In contrast, in 5-minute mating reactions, the *Δbim1* and *Tub1-828-expressing* strains showed 35% and 50% reductions, respectively, in their ability to choose "correct" high expressing partners (Figure 9B).The *Δgim4* mutation did not affect discrimination on its own, nor did it affect discrimination when combined with the *Δbim1* mutation (Figure 9B). Restated, in mating assays dependent on the rapid establishment of a signaling site, loss of Bim1 protein and *Tub1-828-expression* increased variability in transmitted signal η^2^(P) (where P is the ratio P_PRM1_-XFP/ P_ACT1_-YFP, as usual) and impair mating partner choice, while loss of Gim4 only increased variation in η^2^(P).

## Discussion

Although much is known about how cell signaling systems operate, little is known about cell-to-cell variation in their operation. Here we investigated a possible microtubule-dependent process, identified by a genetic screen, that reduces variation. Our results showed that disruptions of microtubule function that affect the nucleus-to-signaling-site bridge increased variation in signal transmission, decreased coherence in population signaling and gene expression, and decreased accuracy of polarity decisions. The loss of coherence in signal and response was entirely suppressed by bypassing the natural means of recruitment of the MAPK scaffold Ste5 to the plasma membrane. Genetic epistasis experiments and direct observation suggested that the microtubule perturbations affected membrane residence of the scaffold protein Ste5.

We first discuss genetic implications of these results and then discuss what they might tell us about mechanism. To test the effects of perturbations on microtubule function, our overall approach (used for some time) (Brent and Ptashne, 1984, Brent and Ptashne, 1985, Colman-Lerner et al., 2005) was to compare quantitative behaviors in different clonal populations of cells of otherwise-isogenic strains with defined genetic differences. We found that reduction of variation required the genes Bim1 and GIM4, which affect cytoplasmic microtubule function, and that it also required other functions, including plus-end attachment to the signaling site, and the ability of the microtubule plus ends to exert force by polymerizing and depolymerizing while attached to the signaling site. Perturbations affecting these functions increased variability in transmitted signal among cells in the population. Moreover, a number of them caused signaling within individual cells to become erratic.

Our work defines *BIM1* and *GIM4* as canalizing genes, affecting stochastic variation around the mean values of measures of signal transmission, independently from their effects on the mean itself. Recent studies have identified a number of genes and alleles that affect variation, rather than mean, in quantitative traits as varied as plant height, flowering and leaf number, intracellular abundances of particular proteins and mRNAs, and the number of somatic cells present in fresh cow milk (Ansel et al., 2008; Fraser and Schadt, 2010; Hall et al., 2007; Jimenez-Gomez et al., 2011; Landers and Stapleton, 2014; Makumburage and Stapleton, 2011; Ordas et al., 2008) (Mulder et al., 2013). 20th century population genetic formalisms (e.g., Hartl (1980)) attributed variation in quantitative traits either to differences in genetic background or to differences in "environment", and a good deal of initial work focused on genetic and other mechanisms by which organisms canalize (or suppress) (Schmalhausen (1949), and by Waddington (1942)) variation due to these two causes. Researchers later recognized a third source of variation, attributed to stochastic events (Ansel et al., 2008; Jimenez-Gomez et al., 2011; Makumburage and Stapleton, 2011) or to unknowable differences in the "microenvironment" (Fraser and Schadt, 2010; Mulder et al., 2013)), for example during development (Waddington, 1957), defined operationally by differences in quantitative phenotype among genetically identical organisms (animals) raised in homogeneous environments (Gärtner, 1990). Some theoretical arguments suggest that genes that canalize traits in the face of genetic and environmental differences (Meiklejohn and Hartl, 2002), and in the face of stochastic differences (Lehner, 2010) might be the same. However, we note that loci that canalize variation in quantitative phenotype caused by differences in environment in other organisms were found by mapping of QTL (Quantitative Trait Loci), so the fact that those loci also reduce variation due to differences in genetic background is not surprising. Moreover, because canalizing loci found by QTL mapping suppressed variation among organisms grown in the same environments, they by definition diminished variation due to "microenvironmental" and stochastic differences. Thus, the fact that BIM1, GIM4, and some canalizing genes affect variation due to all three causes is not unexpected.

Our identification of *BIM1* and *GIM4* as canalizing genes illustrates the frailty of generalizations advanced about such genes based only on population-genetic arguments. These canalizing genes do not operate in some genetic backgrounds (for example, in cells with mutations in proteins that attach microtubules to the signaling site) and would not operate in some environments (for example, those containing chemical inhibitors of microtubule formation). Moreover, in our own previous work, we identified other processes and molecular mechanisms, depending on specific negative feedbacks and specific protein kinases (Colman-Lerner et al., 2005; Yu et al., 2008) that reduced variation in isogenic populations in the same environment, that would not operate in genetic backgrounds deficient in these components. Similarly, recent studies in mice, *Arabodopsis*, and maize reveal canalizing loci, that suppress stochastic variation around mean values, that operate in some environments but not others (Fraser and Schadt, 2010; Jimenez-Gomez et al., 2011; Makumburage and Stapleton, 2011). Thus, taken with previous work, our results establish that genes that reduce variation in specific traits caused by differences in genetic background, differences in environment, and by stochastic causes are not always the same.

In this work, standard deviation, and the related CV and normalized variation used in our previous work (Colman-Lerner 2005, Yu 2008, Pesce 1), provided limited descriptions of variation phenotypes. By eye, measurements of signal transmission over time revealed that clonal populations with some microtubule perturbations had more cells with near average behaviors and a few outliers, while behaviors of cells with other microtubule perturbations were more broadly distributed. Such differences are probably consequential, for example, during metazoan embryonic development and adult tissue maintenance when malfunctioning canalizing processes that increased the number of rare cells that divide or divide out of plane lead to defects and dysplasias. And yet, these sorts of differences in distribution shape were also not well captured by kurtosis, a standard measure of distribution shape that works best with large samples of outlier-free data. We therefore devised a robust and hypotheses-free measure we called "Median Progressive Spread" or MPS, and used it to quantify differences in the distributions of single cell behaviors in mutant populations. This measure worked well for our purposes, and reinforced the point that measures of "deviations from normality" may in some circumstances provide more biological insight into variation than the standard deviation and CV commonly used by population geneticists (Lewontin, 1966).

To understand the mechanism by which microtubule function suppressed variation in signal, we first considered studies by (Maeder et al., 2007, Colon et al. 2016) which reported a gradient of the Fus3 protein kinase originating at the signaling site. We tested a mechanical model in which the microtubule bridge regulates signal strength by precisely positioning the nucleus within the gradient of Fus3, and for which perturbations to microtubule function resulted in a more variable position of the nucleus within the gradient. The underlying assumption in this model wasis that the strength of the signal that reaches the nucleus is stronger when the nucleus is closer to the signaling site. We found no correlation between nuclear distance from signaling site and O, which we used as a proxy for P (masking of correlations between nuclear position and P would require a very specific and unlikely relationship between O and nuclear distance, hence O is a good proxy for P). Our results did not support this simple "motion-in-a-gradient" model, although they do not exclude more complex possibilities, for example that the amount of Fus3 along the gradient is not proportional to signal strength, or that the Fus3 gradient varies so much among individual cells that the distance of nucleus from signaling site is not a good measure of signal level.

We therefore pursued a general approach to learn where in the signaling pathway microtubule-dependent processes reduced variation in signal. To do so, we treated variability in transmitted signal η^2^(P), as a simple (albeit difficult to measure) quantitative trait. We then used the estrogen-inducible synthesis of proteins that activated the PRS ectopically at different steps downstream of the receptor to bypass the pathway steps at which the microtubule perturbations acted to produce the trait. These "suppression by epistasis" (Hodgkin, 2005) experiments showed that the tested microtubule perturbations introduced variation into the signal at steps between Ste5 recruitment and activation of the MAP kinase cascade. They suggested that microtubule function(s) stabilized signaling by membrane-localized Fus3 MAP kinase. We confirmed by direct experiment that enhancement of the variation in signal by microtubule plus-end perturbations at these steps required the Fus3 MAP Kinase. This series of experiments thus extends and generalizes the scope of a classical genetic method first used in the 1930s (Beadle and Ephrussi 1936), and subsequently used extensively in analysis of regulatory pathways in *C. elegans* and *Drosophila* (Avery and Wasserman 1992).

Our epistasis suppression results were consistent with the idea that the microtubule perturbations might interfere with signaling by the MAP kinase Fus3 when localized to the membrane via the membrane-recruited Ste5 scaffold. Ste5 signaling complexes form at the signaling site, which is co-territorial on the membrane with the polarity patch, which defines the site of future polarized growth. We showed that, in some cells in the population, microtubule perturbations slowed the initial establishment and shortened the lifetime of the Ste5 signaling site. We next showed that plus-end microtubule perturbations interfered with a fate decision that depends on correct polarity determination; they interfered with the ability to choose high pheromone-expressing partners in "speed mating" experiments, but not with the ability to choose correct partners given more time. Restated, these experiments showed that perturbations to microtubule function destabilized the signaling site and interfered with rapid polarity decisions.

Taken together, our results suggest a possible explanation for how perturbations to cytoplasmic microtubule function could increase signaling variation and interfere with polarity determination. This idea is based on the fact that the signaling site and the polarity patch both stimulate one another's activity and compete for a limited supply of shared protein components (see Introduction). In this view, unattached microtubule plus-ends disrupt movement and membrane recruitment of proteins, including Cdc42, Cdc20, and Ste5, in their neighborhood. Variable recruitment of Ste5 causes variability in Ste5-dependent signaling, which is then subsequently amplified by the complex positive feedbacks that affect recruitment, trafficking, accumulation, and activation of the numerous proteins and mRNAs, including Ste5 itself, to the signaling site and polarity patch. Moreover, if any of the proteins needed for signaling and polarity determination are transported along microtubules, the fact that the untethered plus-ends cannot hand off cargo to actin cables growing from the signaling site would result in further variability in accretion of signaling and polarity proteins. It is interesting in this context that another protein identified in the high throughput screen, Erv46, is a component of COPII transport vesicles. Restated, in this view, increased variability in Ste5 recruitment to the signaling site causes variability in the amount of signal the site sends via Fus3, and the positive feedbacks amplify this variability. This picture is consistent with our finding that Fus3 is needed for signal variability, and of course with If so, this picture is consistent with conclusions from control theory, in which control of servomechanisms and electronics by positive feedback alone is prone to instabilities.

The complexity of the above model drives home the point that our current means for approaches to perturbing the system and monitoring its operation are not sufficient to elucidate the process under study: how the normal operation of cytoplasmic microtubules helps the cell transmit a signal of constant strength. Too much remains unknown. At the signaling site, there are too many different proteins operating, too many positive cross-regulatory interactions, too many simultaneously occurring mechanical processes like cargo delivery and membrane fusion that are now insufficiently understood. It is as if we had learned to monitor frequency and timing of sounds given off by an electric motor that was operating abnormally because we had disrupted the operation of particular bearings, bushings, and shafts. In this light, our analysis of signal control has explored an "inverse problem", for which inferences from doable experiments are insufficient to describe the system under investigation (Brenner, 2010). But the sort of genetics-powered quantitative physiology we have employed here does have utility. Certainly, it can of course identify important proteins. Moreover, in some cases, it can establish important physical and regulatory processes for which the theory of how human-built machines (eg, Meiksen, 1959, Bennett, 1979) are controlled (here, how positive feedbacks lead to instabilities) can provide small additional insights. Moreover, consider the motor analogy above. In it, different bands of frequencies of noises coming from the partly-malfunctioning machinery will reflect different aspects of motor function. In the same way, here and in the companion paper, we assert that attention to different kinds of system malfunction will lay groundwork for future insight by identifying different aspects or axes of system function dependent on different subsets of system proteins and molecular events.

Reduction of variation by cytoplasmic microtubule function may also be important in metazoans, for coherent population fate decisions (eg proliferation, apoptosis, and differentiation) and polarity decisions needed to form and maintain correct tissue architecture. Cytoplasmic microtubule function in vertebrates is even more complex than that in budding yeast. At the plus-end, mammalian orthologs of yeast Bim1, MAPRE1-3/EB1-3 (Su and Qi, 2001), interact with +TIP proteins, many of which are involved in cell polarity decisions (Sayas and Avila, 2014). These include APC (Adenoma Polyposis Coli), required for radial glial cells to polarize in response to extracellular cues, and for them to support birth and migration of the neurons that make up the cell layers of the cortex (Yokota et al., 2009). Microtubule-end-dependent polarity decisions during adult cell growth and proliferation are important for maintenance of a cancer free soma. APC is a tumor suppressor, frequently inactivated in colorectal cancers (Kinzler and Vogelstein, 1996) and other epithelial tumors (Vogelstein et al., 2013). APC is a negative regulator of the Wnt/β-catenin signaling pathway (Bienz and Clevers, 2000). But APC also carries out other functions suggested by its protein interaction partners (Brent and Finley, 1997), which include plus-end-associating kinesins, formins and actins found at sites of polarized growth (Lesko et al., 2014). At the minus end, lesions in proteins that connect microtubules to the MTOC in the nuclear membrane (in particular the Nesprin-1 and Nesprin-2 isoforms encoded by SYNE1 and SYNE2 (reviewed by Gundersen and Worman (Gundersen and Worman, 2013)) contribute to formation of some solid tumors (eg. squamous cell carcinomas of the head and neck (Stransky et al., 2011)) in which tumor development requires incorrect polarity decisions that allow cells to divide out of an epithelial plane. Human exome sequencing reveals considerable coding sequence polymorphism in genes encoding MAPRE/EB1, APC, and other microtubule-end-interacting proteins in the human population (Fu et al., 2013), suggesting that different allelic forms of these proteins might have quantitative effects on cell polarity decisions in response to signals and so affect cancer incidence (Britain and Brent, unpublished).

#### The microtubule bridge that connects the nucleus to the signaling site

The signaling site and polarity patch are contacted by cytoplasmic or "astral" microtubules. Microtubules are polymers of α/β tubulin heterodimers. Microtubule supply depends on prefoldin, an heterohexameric complex that delivers unfolded proteins to the CCT/TriC chaperonin (Geissler et al., 1998). Prefoldin processes a number of tubular cytoskeletal components, including α-, β-and γ-tubulin. In *Drosophila melanogaster*, mutations in *merry-go-round (mgr)*, which encodes a prefoldin subunit, cause a variety of phenotypes and aberrations in mitosis attributed to incorrectly folded tubulin (Delgehyr et al., 2012). Similarly, yeast mutants lacking prefoldin subunits have reduced levels of α tubulin, and show mitotic delays and malfunctions (Geissler et al., 1998; Lacefield et al., 2006). Two of the yeast components of the prefoldin complex are the proteins Gim4 and Gim2/ Pac10.

Microtubules are attached by their minus ends to protein complexes containing γ-tubulin and accessory proteins, such as Kar1 (Geissler et al., 1998), at structures called microtubule organizing centers (MTOCs). Microtubules elongate and shorten by adding (polymerization) or removing (depolymerization) α/β tubulin heterodimers to their other, plus-ends. Polymerization is blocked by incorporation of dominant negative forms of α-tubulin (for example, in *S. cerevisiae*, one encoded by *Tub1-128* (Anders and Botstein, 2001)). In the nucleus, microtubules form the mitotic spindle, whose orientation sets the plane of mitosis. The orientation of the mitotic plane relative to the cell body is determined by the orientation of the nucleus, which is rotated by a bridge of cytoplasmic microtubules that “anchors" the MTOCs to specific sites in the plasma membrane (Allen and Kropf, 1992). Anchoring at the membrane site requires microtubule plus-end binding proteins that also bind membrane-localized proteins.

In *S. cerevisiase*, the MTOC is a ~1GDa protein structure called the spindle pole body (SPB). It spans the nuclear membrane and binds the minus ends of microtubules on both nuclear and cytosolic surfaces. During budding, which occurs during normal growth, plus-ends anchor at the membrane site of polarized growth. This anchoring aligns the SPB, and thus the longitudinal axis of the pre-division nucleus, with the bud-mother axis (Ten Hoopen et al., 2012).

In pheromone-stimulated cells, cytoplasmic microtubules anchor at the signaling site/polarity patch (Figures 1B and 1E). Thus, anchored microtubules connect bridge the patch and the nucleus. These bridge microtubules alternatively “pull" and “push" the nucleus, which, as a consequence, oscillates within a small volume at the base of the pheromone-induced mating projection, the so-called shmoo tip (Maddox et al., 2003). The pushing depends on Bim1, a plus-end binding protein, that binds only the ends of polymerizing microtubules and anchors them to the membrane. The pulling depends on the Kar3/Cik1 dimer, another plus-end binding complex, which, powered by ATP hydrolysis, anchors and actively depolymerizes microtubules. Bim1-anchored microtubules elongate and push the nucleus away from the site of cell growth, while Kar3/Cik1 anchored microtubules shrink and therefore pull in the nucleus towards the shmoo tip (Zaichick et al., 2009). In pheromone-stimulated *Δbim1* or *Δkar3* mutants, the number of cells with a bridge between the nucleus and the shmoo tip is reduced from 99% to 35-65% (Maddox et al., 2003).

## Methods

General methods, yeast strains and plasmid construction are detailed in Supplementary Information.

### Analysis of cell-to-cell variation in signal and response

We obtained cell images and fluorescent signals as described previously (Colman-Lerner et al., 2005; Gordon et al., 2007).We sonicated cell cultures in microfuge tubes and incubated for 3 hours at 30ºC in medium with 10 μm 1-NM-PP1 and 0, 0.6 nM or 20 nM α-factor. We manually curated timecourses to eliminate cell outlines showing segmentation mistakes, overlapping cells and other imaging artifacts. In these assays, cell densities were in the OD_600_ 0.02 – 0.1 range.

To analyze the data, as in Colman-Lerner et al. 2005, we bounded the system and defined total system output by the total accumulated signal from a fluorescence protein reporter gene driven by a pheromone-induced promoter (*PRM1*). We then divided the PRS into a "pathway subsystem", which included all the events leading to activation of transcription, and an “expression subsystem”, which includes all the events from transcription initiation through accumulation of expressed protein. In this framework, the system output for any given cell O_i_ was the product of the average pathway subsystem output per unit time *P_i_*, which varies with input pheromone dose, the expression subsystem output *E_i_*, and the duration of stimulation ΔT (Colman-Lerner et al., 2005), as follows:

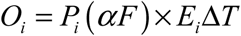

We considered output of each subsystem to be the sum of the capacity of the subsystem in each cell (i.e., "capacity") plus stochastic fluctuations in the operation of that subsystem during the course of an experiment (“noise”): *L* and λ, respectively, for the pathway subsystem, and *G* and γ for the expression subsystem. Thus,

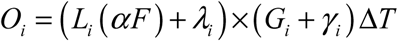

We defined the variation in system output as the normalized variance of in *O_i_*, η^2^(O) decomposable into the sum of individual sources and a correlation term (Colman-Lerner et al., 2005), as follows,

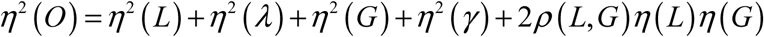

In the WT, in the deletion strains, and mutants used in the manuscript, we measured output of each reporter; as well as η^2^(P) (i.e. η^2^(L) + η^2^(λ)), pathway variation, for the pheromone response system. We estimated η^2^(P) as the variance in the difference of the normalized abundances of the two fluorescent proteins, one driven by the P_PRM1_ promoter and the other driven by the constitutive, pheromone-independent promoter (P_ACT1_ or P_BMH2_, depending on the strain). This variance is actually equal to η^2^(P) + η^2^(γ) (see supplement), but we have previously shown that η^2^(γ) is small enough (Colman-Lerner et al., 2005) to make our estimate a good approximation for η^2^(P).

To quantify the perception that in microscopic time course measurements, the spread of trajectories for cells of some strains differed in both magnitude and quality from the corresponding spread for other strains, we developed a statistical measure we called the progressive spread distribution (PSD). For each strain, we computed the PSD by first computing the absolute deviation from the median of each cell's final pathway output. We then sorted these deviations from smallest to largest. We found the first value of the PSD by starting with the two trajectories with smallest absolute deviations and finding the difference between the corresponding pathway output values. We computed the next value of progressive spread by including the value of pathway output corresponding to the cell trajectory with the third smallest deviation, and again finding the range of corresponding outputs, and so on until we had included trajectories for all the cells in each tested population of each strain. For a given set of pathway output measurements, the last value of the PSD was simply the full range of the pathway output values. For symmetric distributions, the median of the PSD, which we call Median Progressive Spread (or MPS), is equivalent to the interquartile range.

### Chemical inhibition of microtubule function

For every experiment in every strain, we grew cells in medium with microtubule-polymerization inhibitors for 3 hours, then counted % cells arrested in metaphase. If the proportion of arrested cells was >95%, we declared that the treatment had worked. For the experiments in the GPY4000 reference strain, we also made a derivative that expressed GFP-Tub1 instead of Tub1 and observed disappearance of visible microtubule bundles after chemical inhibition.

### Analysis of nuclear movements

We defined nuclear position in two ways. One was by the position of the SPB, which attaches microtubules to the nucleus. To do so we used otherwise-isogenic cells that expressed an SPB component, Spc42, tagged with Green Fluorescent Protein. We defined the signaling site using RFP-tagged Sec8, verifying its localization to the shmoo tip by inspection of brightfield images (not shown). We divided cells into two aliquots. We treated one with 50 nM pheromone, a concentration at which most receptors are occupied, cells shmoo, and there is low cell-to-cell variation in transmitted signal (low η^2^(P)) (Colman-Lerner et al.2005). We treated the other aliquot with 1 nM pheromone (the EC 50 dose), a concentration that half-saturates receptor and arrests progression through the cell cycle. At this concentration cells do not shmoo, but form a broad mating projection known as a peanut (Figure 3A iii) and show high cell-to-cell variation in transmitted signal (*η^2^(P)*). The other means to track nuclear position was to mark the nuclear volume with a fluorescent signal. To do so, we used strains expressing HTB2-YFP, a histone subunit, and used the *P_PRM1_-mCherry* and *P_ACT1_-CFP* reporters to measure pathway output in the same cells.

### Analysis of Ste5 membrane localization

Ste5 protein is present at very low numbers. There are ~500 molecules per cell on average (Thomson et al., 2011), and due to significant YFP fluorophore maturation times, when Ste5 is tagged with YFP only ~350 of 500 YFP-Ste5 are fluorescent in exponentially growing cells (Gordon et al.). In these cells, the fluorescence signal from YFP-Ste5 molecules is barely above background autofluorescence. To increase the signal-to-background ratio, we used strain MW003 (Ventura et al., 2014), a strain that expresses three copies of chromosomally integrated *STE5-YFP(x3)*, in which three copies of YFP are fused to 3' end of the *STE5*, expressed from native promoter sequences. For this fusion protein, tripling the gene dosage increases the signal-to-noise ratio while only modestly perturbing levels of Ste5 and system activity, as reflected by the amount of reporter gene output (about 1.5 times wild-type cells, data not shown). We stimulated this strain's cells with 1 μm pheromone (high pheromone) collected scanning confocal fluorescent microscope images every 10 minutes over 4 hours (see Supplement).

To quantify Ste5 patches, we created image montages of each cell automatically using Rcell´s function *cimage.* These montages included all Z slices for every time point in an ordered layout. We identified Ste5 patches manually and tagged them using ImageJ's plugin Cell Counter on the image montages. Cells were presented in random order (with no identifying label) to avoid any bias in the manual quantification. The tag data created with Cell Counter was then mapped to the corresponding cell in the Cell-ID dataset (Bush et al., 2012; Gordon et al., 2007) for further analysis in R.

We assessed the significance of differences in quantitative variables (e.g. time to 1^st^ pol, 1^st^ pol duration, etc.) using permutation tests (Good, 2004), which gave results consistent with traditional parametric tests (e.g. t-test, F-test of variance, etc). We assessed the significance of differences in the fraction of cells with a given phenotype (cells with coexisting polarizations for more than 60 min, cells with no polarizations, etc.) using Fisher´s exact test for count data. We calculated standard errors for all variables using bootstrap resampling (Efron, 1979).

### Partner choice mating assays

The rapid mating protocol is derived from a previously published method (Jackson and Hartwell, 1990). We grew MATa and MATalpha cells overnight to exponential phase. In the morning we counted the number of cells/ml in each culture with a hemocytometer. Next, we made a 50:50 mixture of GPY4077 (Reference or "WT" MATalpha *Δade2*) and GPY1551 (MATalpha *Δalpha1 ADE2+*) cells. We added approximately 10^7^ cells of this mixture (about 1 ml) to 15ml BD Falcon tubes. To supplement the concentrations calculated from the hemocytometer, we counted plated dilution series from the MAT alpha mixture on YPD plates in quadruplicate. On these plates, each viable GPY4077 cell (or cell clump) will form a pink colony and each viable GPY1551 cell (or cell clump) will form a beige colony. We sonicated the cultures of reference MAT<u>a</u> cells (GPY4078) and otherwise-isogenic *Δbim1, Δgim4, Δbim Δgim4*, and *Tub1-128-expressing* perturbed cells to be tested, counted them in a hemocytometer, and plated out dilutions as above.

We performed all mating experiments in triplicate. To carry out the mating, we added 10^6^ MATa cells (about 0.1ml) to the 10^7^ cells in the MATalpha mix Falcon tube, vortexed, and vacuum filtered the cells onto a 0.2 μm Millipore nitrocellulose filter. We placed the filter face up on uracil-deficient agar medium and incubated at 30ºC for 5 minutes. We transferred the filters to 5 ml of SD-N (nitrogen deficient medium; H_2_O, 2% glucose, and 1X YNB with no ammonium sulfate or amino acids) and vortexed for 30 seconds at maximum power with a Fisher Vortex Genie 2. We sonicated these mating mixtures for three minutes at power 3, and plated ~200 cells from the mating mixtures (having diluted cells to densities estimated from previous hemocytometer counts) onto SD deficient in lysine, methionine, and adenine using 3mm glass beads. On this medium, only diploids grow. On this medium, diploids resulting from mating of MATa GPY4078 cells with "correct" MATα wild type GPY 4077 cells form pink colonies, diploids resulting from mating with lower-pheromone-expressing GPY1551 strain form white colonies. We scored the plates by hand after two days of incubation at 30°C. In control experiments against GPY4007 cells, under these conditions 3-5% of reference MATa cells and *Δbim1* and *Tub1-828-expressing* cells mated and formed diploids.

## Author contributions

GP and SZ performed the microtubule perturbation and epistasis suppression experiments. WP showed the Fus3 dependence of the introduced variation and analyzed, plotted, and determined statistical significance of quantitative results. AB and VR quantified effects of microtubule perturbations on the Ste5 patch at the signaling site. RCY contributed to experimental design, data analysis and earlier versions of the manuscript. RB, GP and AC-L directed the work and its interpretation, wrote the paper, and guarantee the integrity of its results.

## Acknowledgments

We are grateful to Steve Andrews, Alexander Mendenhall, and Bryan Sands for valuable discussions throughout. Work benefited from ability to use SGD and contacts with SGD staff particularly Michael Cherry and Stacia Engel. Work was supported by R01 GM097479 to RB. Earlier work received support from grants R01 GM086615 to RB and RCY and P50 HG002370 to RB.

## 1 General methods

We performed DNA manipulations including PCR and subcloning as described (Ausubel et al., 1987–Ausubel et al., 2016). We cultured and manipulated yeast as described (Ausubel et al., 1987–Ausubel et al., 2016; Guthrie and Fink, 1991). Unless otherwise noted, we grew cells in synthetic dextrose complete (SDC) media consisting of Brent Supplemental Media (MP Biomedicals, Solon, OH), yeast nitrogen base without amino acids and ammonium sulfate (BD, Franklin Lakes, NJ), and dextrose (Sigma-Aldrich, St Louis, MO).

## 2 Yeast strains

### 2.1 *Base strains*

#### *2.1.1* Construction of GPY1802, a strain derived from BY4741

We first generated GPY1802, a BY4741 derivative with markerless *Δbar1* and *cdc28-as2* loci. To do this we first deleted *BAR1* in BY4741 using a PCR product containing the pRS406 URA3 gene flanked by 50 nt of homology the *BAR1* promoter and terminator. We confirmed proper insertion was confirmed by PCR and a phenotypic test of *BAR1* status. We named the resulting *Δbar1::URA3* strain GPY8190.

We then deleted the *URA3* marker by transforming GPY8190 with a 100-bp double stranded oligonucleotide composed of two "arms" of 50 nt of homology to the sequences flanking the URA3 marker. We selected for *Δura3* transformants in 5-FOA plates and confirmed proper deletion by PCR. We named the resulting strain GPY1800 and annotated this locus in the strain table as *Δbar1* (Table S4).

Next we introduced the *cdc28-as2* allele. We used linearized pCDC28-as2-406 to "loop in" the allele, generating strain GPY1801, and then we "looped it out" as described earlier (Colman-Lerner et al., 2005). We confirmed proper recombination by PCR and confirmed it by testing for 1-NM-PP1 sensitivity. We named the resulting strain GPY1802 and annotated this marker in the strain table as *cdc28-F88A* (Table S4).

##### Construction of GPY1804, as GPY1802 but without a constitutive reporter

We used GPY1802 to generate the *TUB1-828-expressing* strain and its cognate reference strain (see below). To do so, we added a P_PRM1_-mCherry reporter to GPY1802 by replacing the PRM1 ORF with the mCherry coding sequence followed by a terminator and a marker. We used an approach identical to the one above for P_PRM1_-YFP and mRFP, except that in this case we used an mCherry-T_ADH1_---G418 r template described before (Yu et al., 2008). We confirmed integration by PCR and checked the presence of a pheromone-inducible mCherry reporter. We named the resulting strain GPY1804.and annotated this locus as *PRM1pr-mCherry-G418(MX6)* in the strain table (Table S4).

##### Construction of GPY 4000-derived strains to validate and further analyze Δbim1 and Δgim4 phenotypes

In other experiments, we had learned that *P_ACT1_* was mildly induced by pheromone. For GPY4000 and related strains, we therefore chose to use *P_BMH2_-YFP* as a constitutive promoter to ensure that the changes in η^2^(P) observed in the deletion mutants did not arise from a change in variation in the pathway controlling the constitutive control reporter.

We introduced all gene deletions in the GPY4000 strain by a PCR approach. We first obtained strains carrying the gene deletion of interest marked with G418 resistance from the haploid deletion collection. We then changed the G418 resistance to either NAT resistance or hygromycin B resistance by transforming the deletion strain with a PCR product of the resistance cassettes and selecting for the marker carried in the PCR product. Since all three drug resistance markers contain the same 300-400 bp promoter and terminator, those flanking sequences directed a double homologous recombination gene replacement. Proper recombination was confirmed by the loss of the original G418 resistance. Subsequently, we amplified the gene deletion locus by PCR, including 300-500 bp of 5' and 3' gene-specific flanking sequences in the amplicon. We transformed this PCR product into GPY4000, selected for the appropriate drug resistance and confirmed proper gene deletion by PCR. The resulting strains are listed in Table S1.

##### Construction of SGA101, a clean background for testing additional genes that might affect microtubule function

SGA85 was is the haploid MATa reporter strain used in the companion paper (Pesce et al. 2016) as a reference in our screen to find otherwise isogenic deletion strains with alterations in variation in signal transmission. In this study, we used this strain to introduce different alleles possibly affecting microtubule function by loop in/ loop out replacement with *URA3*-marked plasmids and for the three-color FP live cell time courses (see below). Starting with the reference strain SGA85, we derived a *Δura3* strain by transforming SGA85 with a PCR product spanning the entire *LYP1* locus from promoter to terminator, amplified from BY4741 genomic DNA. We selected for loss of the *URA3* marker in 5-FOA plates and confirmed restoration of the *LYP1* ORF by verifying the loss of YFP fluorescence from P_ACT1_-YFP-T_ADH1_ and by diagnostic PCR. The resulting strain was a *URA3 LYP1* derivative of SGA85. We named it SGA101.

### 2.2 *Construction of strains with genetic alterations in KAR1, KAR3, and CIK1*

#### kar1-Δ15 (SGA109).

We obtained plasmid pMR1593 as a gift from Mark Rose and used it as described (Vallen et al., 1992) to replace *KAR1* for *kar1-Δ15* in parent strain SGA101. Briefly, we linearized pMR1593 with Bgl II, transformed it into SGA101, selected for uracil prototrophy and in a second step selected for loss of the *URA3* marker in 5-FOA plates. We screened the resulting colonies by PCR to identify a *kar1-Δ15* strain, which was named SGA107. For this strain and the kar3-1 (SGA108) strains and the *Δkar3 (GPY4003)* strain, the reference strain is SGA103.

We then transformed SGA107 with pTC-PBMH2-YFP- TADH1-URA3 linearized with StuI as above. We identified transformants with a single PBMH2-YFP copy using fluorescence microscopy. The resulting strain was named SGA109.

#### kar3-1 (SGA108)

We obtained the plasmid pMR1510 from Mark Rose and used it as described to replace *KAR3* with *kar3-1*. Briefly, we linearized pMR1510 with MluI, transformed it into SGA101, selected for uracil prototrophy and in a second step selected for loss of the *URA3* marker in 5-FOA plates. We screened the resulting colonies by PCR to identify a *kar3-1* strain, which was named SGA106. As above, we transformed SGA107 with linearized pTC-PBMH2-YFP-TADH1-URA3 and identified transformants with a single PBMH2-YFP copy using fluorescence microscopy. The resulting strain was named SGA108.

#### Δkar3 (GPY4003)

To make strains with a deletion in the KAR3 coding sequence, we used primers GP800-801 to amplify the hygromycin B resistance cassette hph from pAG32. This PCR was transformed into GPY4000, which was then plated on non-selective media and replica plated to media containing hygromycin B. Transformants were confirmed to be knockouts of KAR3 using primers GP1397-1044, and the resulting strain was named GPY4003.

#### Δcik1 (GPY4123)

To make strains with a deletion in the CIK1 coding sequence, primers GP1361-1362 were used on KAN MX4 to generate a PCR product that had the TEFpr-G418-TEFter cassette with homologous tails directly flanking the 5' and 3' of the CIK1 ORF. GPY4104 was then transformed with the generated PCR product, plated on non-selective media, and replica plated to media containing geneticin. Transformants were confirmed to be knockouts of CIK1 using primers GP1397-1062, the resulting strain was named GPY4123. The reference strain for this Cik1 strain is GPY4000 4741 MATa Δbar1::Δmarker cdc28-F88A Δprm1::PRM1pr-mCherry--G418(MX6) BMH2::BMH2pr-YFP--MET15

### 2.3 *Construction of strains with nuclear label and gene expression reporters for nuclear movement experiments*

#### HTB2-YFP strains

We based this set of strains was based on SGA85, the reference strain for the modified haploid gene-deletion collection used in our screen for cell to cell variability mutants. To make it, we modified SGA85 by removing the P_PRM1_-CFP-T_ADH1_ and P_ACT1_-YFP reporters and introducing P_ACT1_-CFP reporter and a YFP label to allow visualization of the nucleus.

In SGA85, the P_ACT1_-YFP reporter had been inserted in the *Δlyp1* locus replacing the LYP1 ORF, as part of an inserted plasmid with a URA3 marker. We removed the entire plasmid insertion by transforming SGA85 with a PCR product of the WT LYP1 locus that included 300-500 bp of promoter and terminator sequences. We selected for ura3 mutants in 5-FOA plates and identified the colonies with a restored LYP1 locus by their loss of YFP fluorescence, acquired thialysine sensitivity and by PCR. This LYP1 variant of SGA85 was named SGA101.

In SGA85, the *P_PRM1_-CFP* reporter had been inserted in the *Δbar1* locus replacing the BAR1 ORF, as part of an inserted plasmid with a HIS3 marker. We removed the entire plasmid insertion by transforming SGA85 with a PCR product of a *Δbar1::URA3* locus that included 300 bp of promoter sequences and 700 bp of terminator sequences. We used strain GPY8190 as template for the *Δbar1::URA3* PCR. We selected for uracil prototrophs and confirmed appropriate gene replacement by the loss of the pheromone inducible CFP reporter, acquired histidine auxotrophy and by PCR. We named the resulting strain SGA114.

We then introduced a nuclear-localized fluorescent label by fusing the HTB2 ORF to YFP. We used a PCR product containing the YFP- TADH1-KAN(MX4) cassette with ends with homology to sequences upstream and downstream of the HTB2 ORF, that was generated with the YFP-KAN(MX4) from the Pringle collection (Longtine et al. 1998). We confirmed proper insertion by the appearance of a bright yellow fluorescence label in the nucleus and by PCR. We named the resulting strain SGA118.

We recovered the URA3 marker by transforming SGA118 with a 200 bp double stranded oligonucleotide composed of two 100 bp sequences with homology to the ends of the URA3 marker in the Δbar1::URA3 locus. We plated the transformation in 5-FOA and checked proper deletion of URA3 by PCR. We named the resulting strain SGA123.

We next deleted BIM1 from SGA123. We followed the same protocol described above for deleting BIM1, except that we used HIS3(MX4) as the selective marker. The resulting *Δbim1* strain was named SGA125.

#### SEC8-RFP SPC42-GFP strains

This set of strains was derived from a clonal isolate from the yeast GFP collection (Huh et al., 2003). The members of this collection were derived from a BY4741 parental strain. The clone we used carried a modified allele of *SPC42* expressing a Spc42 with a C-terminal GFP tag and linked to a *HIS3* marker (this allele is denoted here as *SPC42-GFP—HIS3).*

To make the reference strain for this set, we first deleted the *BAR1* gene in *SPC42-GFP— HIS3* using the approach we described above for the generation of GPY8190.

This resulted in strain GPY1752: *SPC42-GFP—HIS3 Δbar1::Δmarker.*

In a second step, we modified the *SEC8* gene to express Sec8 with a C-terminal mCherry tag. We used an approach identical to the one above for P_PRM1_-YFP and mRFP, except that in this case we added the tag at the end of the gene instead of replacing the gene with it, and used the mCherry-T_ADH1_---G418 r template described before.

This resulted in strain *SPC42-GFP—HIS3 SEC8-mCherry—KanMX6 Δbar1::Δmarker*, named GPY1751.

In a final step to construct the base strain for this series, we replaced the *CDC28* gene by the analog-sensitive *cdc28-F88A* allele. We used linearized pCDC28-as2-406 to "loop in" the allele, and then we "looped it out", as described earlier (Colman-Lerner et al., 2005). We confirmed proper recombination by PCR and confirmed it by testing for 1-NM-PP1 sensitivity.

This resulted in strain *SPC42-GFP—HIS3 SEC8-mCherry—KanMX6 Δbar1::Δmarker cdc28-F88A*, named GPY1752. GPY1752 was the base strain for this set.

Starting with GPY1752, we generated GPY1709 *(Δbim1--NatMX6).* We used GPY1752 and GPY1709 to measure the effect that deleting *BIM1* had on nuclear positioning relative to the site of polarized growth (Figure 3).

To measure the effect that deleting *GIM4* had on nuclear positioning we used otherwise-isogenic strains that carried GFP-tagged alpha tubulin, in addition to Spc42-GFP and Sec8-mCherry. We to make these, we transformed GPY1752 with plasmid pAFS125C, a construct from Aaron Straight carrying the *TUB1* gene, driven by its own promoter and fused in-frame with a gene coding for GFP (Straight et al., 1998). pAFS125C is an integrative plasmid (has no yeast origin of replication) and is marked with the *URA3* gene. Before transformation we linearized pAFS125C ( a gift of by Julia Kardon, a postdoc in Tania Baker's lab at MIT). cutting the *TUB1* promoter with the Eag I restriction enzyme. This cut site created a strong preference for insertion of the plasmid at the *TUB1* promoter in the genome. The resulting *TUB1* locus carried two tandem copies of the *TUB1* gene. The 5' copy was driven by the endogenous *TUB1* promoter and fused with the GFP gene. The 3' copy was driven by the plasmid-borne *TUB1* promoter and is not tagged. We noted this modified *TUB1* locus as *TUB1::PTUB1-GFP-TUB1---URA3.*

We named the resulting strain GPY1756 (a derivative of GPY1752 carrying *TUB1::PTUB1-GFP-TUB1---URA3*). GPY1756 and GPY152 showed no difference in nuclear positioning (data not shown). Starting with GPY1756, we generated GPY1759 (*Δgim4--NatMX6*). We used GPY1759 to measure the effect that deleting *GIM4* had on nuclear positioning relative to the site of polarized growth (Figure 3).

### 2.4 *Construction of strains expressing human estrogen receptor chimeras*

#### Vectors for homologous recombination with excisable CEN/ARS cassettes

For some cloning steps below we used the pTC41x series of derivatives of the pRS41x shuttle vectors (Gordon et al., 2007). Each pTC41x vector was made by cloning a CEN/ARS containing-fragment flanked by AatII overhangs into the AatII site of each pRS40x vector. As a result all pTC41x vectors can be converted to pRS40x versions ("integrative" vectors that lack a yeast origin of replication) by cutting with AatII and religating.

#### Plasmid carrying estrogen receptor chimera driven by the ADH1 promoter

Our source of the GAL4 DNA binding domain – human estrogen receptor – VP16 transactivation domain (GEV) chimera was plasmid pp1557, a gift of Peter Pryciak (Takahashi and Pryciak, 2008). We first subcloned a PvuI fragment of pp1557 containing P_ADH1_-GEV-T_ADH1_ into the SmaI site of pTC415. To do this we co-transformed PvuI-cut pp1557 and SmaI-cut pTC415 in yeast strain BY4741. We isolated gap-repaired plasmids from leucine protrotrophs using a Zymoprep Yeast Plasmid Miniprep I kit. The resulting plasmids were then transformed into Z-competent *E.coli* to amplify the plasmid, 8 colonies were grown overnight in LB-Amp media and miniprepped using Qiagen QIAprep Spin Miniprep Kit, and the plasmid sequences were confirmed by DNA sequencing. This cloning resulted in pSZ100.

We subsequently removed the CEN/ARS in pSZ100 by the AatII process described above, yielding pSZ101. pSZ101 elements are P_ADH1_-GEV-T_ADH1_---LEU2.

#### Plasmid carrying estrogen receptor chimera driven by the BMH2 promoter

In a separate line of research we had found that the *ADH1* promoter showed unusually high cell-to-cell variability. For this reason, in parallel we constructed an alternative GEV expression vector driven by the *BMH2* promoter. We amplified the GEV coding sequence by PCR from pp1557, using primers that added to the amplicon ends with homologies to the 3' end of the *BMH2* promoter (in the 5' end of the amplicon) and to the pTC7 backbone (in the 3' end of the amplicon, pTC7 is the plasmid from which pTC-P_BMH2_-YFP-URA3 is derived (Colman-Lerner et al., 2005)). We cut pTC-P_BMH2_-YFP-T_ADH1_-URA3 with EcoRI (partial digest) and XhoI to remove the YFP sequence. We co-transformed this digested plasmid with the GEV PCR product into yeast strain TCY3277, which carried a P_GAL1_-GFP-T_ADH1_--HIS3 reporter. We transferred to *E. coli*, mapped and sequenced plasmids that showed estradiol induction of GFP.

We named the resulting pTC-P_BMH2_-GEV-T_ADH1_--URA3 construct pSZ110.

Subsequently, we moved the P_BmH2_-GEV-T_ADH1_ cassette to LEU2 vector pRS305. We cut pSZ110 and pRS305 with BglI and ligated both DNA's by standard cloning procedures. Recombinant plasmids were screened for those derived from a pRS305 backbone and containing the P_BMH2_-GEV-T_ADH1_ cassette. The resulting plasmid was named pSZ111.

#### Strains carrying single copies of P_BMH2_-GEV or P_ADH1_-GEV

Both P_BMH2_-GEV or P_ADH1_-GEV LEU2 plasmids are designed to be integrated by single recombination in the *LEU2* locus, such that one or more copies could be integrated at the *LEU2* locus. It was essential for our purposes to ensure that the strains we used had just one copy of either construct.

To obtain strains with single copies of the GEV constructs, we first obtained a set of clones with unknown number of copies for each construct. To determine the number of GEV construct copies integrated in the clones we introduced pp1744 (a gift from Peter Pryciak containing P_GAL1_-GFP-T_ADH1_ ---- His5_s.pombe_), linearized at the *HIS3* locus with Nhe1, into each of them and screened the P_GAl1_-GFP transformants to identify those with just one copy of this GFP reporter. Finally, we stimulated the GEV clones with a gradient of estradiol doses and obtained a GFP vs estradiol dose response. We use the dose responses to identify the clones with just one copy of the GEV constructs. For subsequent applications of these strains, we used the parents of these clones, because they did not carry the P_GAL1_-GFP construct. The steps we followed are described below.

We generated 8 clones carrying P_BMH2_-GEV by transforming pSZ111 linearized with XcmI at the LEU2 locus into GPY1804. 8 leucine protrotophs were saved and named GPY1806 (ah). Similarly, we obtained 12 clones carrying P_ADH1_-GEV by linearizing pSZ101 with a PacI cut within P_ADH1_ and transforming GPY1804. 8 leucine protrotophs were saved and named GPY1805(a-h). We then transformed all 16 of GPY1806(a-h) and GPY1805(a-h) with pp1744 linearized with Nhe1.

We determined how many copies of pp1744 were integrated in 2 isolates of each of the 16 transformations by measuring GFP levels after incubating cells for 3 hours in 2% galactose. We selected 1 clone with one copy of pp1744 for each of the 8 clones each of GPY1806 and GPY1805 we had isolated earlier and named them GPY1808 (a-h) and GPY1807 (a-h) respectively. Finally, we did estradiol dose – response curves in all 16 GEV clones carrying P_GAL1_-GFP. We grew GPY1808 (a-h) and GPY1807 (a-h) in glucose media overnight in exponential phase and stimulated these cells at low density with a gradient of estradiol concentrations (1 to 50 nM range) for 3 hours. After 3 hours we added 200μg/ml cycloheximde, incubated cells 5-9 hours at room temperature and measured GFP levels by flow cytometry. In each group of 8 clones, a majority of 5 to 7 clones showed identical dose responses. The remainder showed higher maximal induction and lower EC50. We judged that the dose responses shown by the majority of the clones corresponded to the single-copy integrants. From this point forward the names GPY1805, GPY1806, GPY1807 and GPY1808 (without the lower case letters sub-ID) were used to designate a single copy GEV integrant of each set.

The GEV locus driven by the BMH2 promoter is annotated as *BMH2::BMH2pr-GAL4BD-hER-VP16–LEU2* in the strain table (Table S1).

#### Comparison of variability of activation dependent on P_BMH2_-GEV or P_ADH1_-GEV constructs

We first integrated a housekeeping fluorescent protein reporter in GPY1805 and GPY1806 by integrating pTC-P_BMH2_-YFP-T_ADH1_—URA3 linearized with StuI within P_BMH2_. We identified strain with single-copy integrations and named them GPY1809 (P_ADH1_) and GPY1810 (P_BMH2_). Then we performed estradiol dose responses as above and analyzed the results as described earlier (Colman-Lerner et al., 2005). From this analysis we found that the level of cell-to-cell variation in GEV activator function (amount and activity) in GPY1809 (P_ADH1_) was higher than in GPY1810 (P_BMH2_).

Based on these studies we decided to use GPY1810 (P_BMH2_) for subsequent constructions. At this point we decided to free the URA3 marker to allow the introduction of the *TUB1-828* construct (see below). We thus replaced the pTC-P_BMH2_-YFP-T_ADH1_—URA3 integrant with a pTC-P_BMH2_-YFP-T_ADH1_—MET15 integrant by loop out using 5-FOA selection followed by transformation with the MET15 plasmid linearized by cutting within P_BMH2_. We screened the transformants by fluorescence microscopy to identify single-copy P_BMH2_-YFP integrants. We called the resulting strain GPY1858. The locus is annotated as *BMH2::BMH2pr-YFP--MET15* in the strain table.

### 2.5 *Construction of an estradiol inducible* TUB1-828-expressing *strain (GPY1873)*

We used an inducible *TUB1-828* construct that we received from Kirk Anders (Anders and Botstein, 2001). This plasmid, pRB2949, is a CEN/ARS URA3 yeast episome carrying a *P_GAL1_-TUB1-828* cassette. Expression of *TUB1-828* from this construct is induced by galactose and repressed by glucose.

In our experimental system, incubation in media without glucose with galactose alters many aspects of system behavior, including a substantial increase in pathway variation (data not shown). We therefore induced *TUB1-828 expression* using the estradiol responsive Gal4-ER-VP16 chimeric transcription factor (described above, Louvion et al., 1993).

We transformed GPY1858 with plasmid pRB2949 (Anders and Botstein, 2001). We tested several colonies for growth in the presence of estradiol. All of them failed to grow, the expected consequence of the induction of *TUB1-828.* We selected one clone for *TUB1-828* expression experiments and named it GPY1873.

After constructing GPY1873 and testing its effect on the PRS response to pheromone we found that basal *TUB1-828 expression* conferred by the estrogen-responsive chimeric transcription factor was enough to give the maximum response observed, shown in Figure 7B.

### 2.6 *Construction of strains expressing ectopic PRS activating proteins for bypass experiments*

If strains that contain artificial activator plasmids are grown on media that is depleted of glucose, the artificial activator induces the pheromone pathway and stops growth. For that reason, when using antibiotic selection to knock out genes, we first re-suspended these strains in 15ml of SDC medium for 6 hours, spun down, re-suspended in 100μl, and plated directly onto antibiotic. This transformation protocol ensured that the cell cycle was not inhibited by expression of the ectopic activators prior to antibiotic selection.

In order to screen for single copy integration of activator plasmids, we used frozen stocks of 6 isolates to inoculate cultures in selective SDC media. These cultures were allowed to grow for 6-8 hours and were then diluted to grow in log phase overnight. The following morning the 6 isolates were induced cultures that contained serial dilutions of β-Estradiol in SDC with .04mg/ml casein and 10μm 1-NM-PP1. These cultures were then fixed in 200μg/ml cycloheximde and taken to the flow cytometer for fluorescence measurements. Out of 6 isolates tested, the 4-5 that typically had the same dynamic range and maximum induction were scored as single integrations and given strain numbers.

#### STE4 activator strains

GPY1810 was transformed with pp1610 linearized with BsmI to target the plasmid to the *HIS3* locus and plated onto histidine deficient media. 6 isolates were then streaked to single colonies, patched, and immediately frozen to prevent unwanted activation of the pheromone pathway. One of the isolates that was determined to have 1 copy of *GAL1pr-STE4* was named GPY1816 and used as base strain in this set. We noted this locus *his3-Δ1::GAL1pr-STE4--HIS3.* This strain was then transformed with the PCR product generated from primers GP1046 and GP1047, mentioned above, to knock out BIM1. 8 isolates were then screened for successful *Bim1* removal using primers GP1246 and GP1397, the positive isolate was named GPY1818. Later, upon deciding to incorporate *Tub1-828* into our studies, we needed to free up *URA3* from these strains. To do so, we plated, GPY1816 and GPY1818 onto 5-FOA and collected singles, 4 of which were screened to verify the successful loop-out of the *BMH2pr-YFP::URA3* plasmid. The resulting strains were named GPY1828 (STE4) and GPY1830 (STE4 *Δbim1)* and transformed with a plasmid containing *BMH2*pr-YFP::*MET15* linearized at the *BMH2* promoter with Stu1. 6 transformants of each were streaked to singles and single copy integrations were determined by microscopy, the resulting strains were named GPY1855 and GPY1862. GPY1855 was then transformed with the PCR product generated using primers GP1153 and GP1155, mentioned above, to knock out *GIM4*, transformants were then PCR verified for the knock out using primers GP1175 and GP1397. GPY1855 was also transformed with a *Tub1-828* CEN/ARS plasmid pRB2948, received from Kirk Anders (Anders and Botstein, 2001), and plated directly onto *URA3* selective plates. Positive transformants were identified by cell cycle arrest in response to 50nM −Estradiol and named GPY1879.

#### STE5-CTM Δste5 strains

GPY1810 was transformed with pp1611 linearized with BssHI to target the plasmid to the *HIS3* locus and plated on histidine deficient media. 6 isolates were then streaked to singles, patched, and immediately frozen to prevent unwanted activation of the pheromone pathway. The isolate that was determined to have 1 copy was named GPY1817. This strain was then transformed with the PCR product generated from primers GP1046 and GP1047, mentioned above, to knock out BIM1. 8 isolates were then screened for successful *Bim1* removal using GP1246 and GP1397, the positive isolate was named GPY1820. The same protocol was followed as for *STE4* to free up the *URA3* marker for the incorporation of *Tub1-828* into GPY1817 and GPY1820. The resulting strains that contained *BMH2*pr-YFP::*MET15* instead of *BMH2*pr-YFP::*URA3* were named GPY1856 (*STE5*-CTM) and GPY1864 (*STE5*-CTM *Δbim1*). GPY1856 and GPY1864 were then transformed with PCR DNA with tails homologous to the *STE5* promoter and terminator generated using primers GP1098 and GP1099 on the NAT MX4 plasmid to knock out *STE5* from the genome. Transformations were grown for 6 hrs in SDC and plated directly to proper selection plates. Transformants were streaked out on selective plates to singles and PCR verified for the deletions using GP1101 and GP1397, positive isolates were named GPY1915 (*STE5*-CTM *Δste5*) and GPY1916 (*STE5*-CTM *Δbim1 Δste5*). GPY1915 was then used to incorporate the *Δgim4* and *TUB1-828* mutations as described for the *STE4* activator to generate GPY1997 and GPY1986 respectively.

#### STE11-4-STE7 Δste5 strains

GPY1806 was first transformed with a ds PCR product that had tails homologous to the *STE5* promoter and terminator, generated using primers GP1098 and GP1099 on the NAT MX4 plasmid. We used this PCR product to knock out *STE5* from the genome. The transformation was first plated on leucine deficient plates, allowed to grow to a lawn overnight, and then replicated to leucine deficient plates with containing NAT. Transformants were streaked out on selective plates to singles and PCR verified for the deletions using primers GP1101 and GP1397. We named one positive isolate GPY1861. GPY1861 was then transformed with *BMH2*pr-YFP::*MET15* and single copy isolates were identified as described for the *STE4* activator to generate GPY1874. GPY1874 was then transformed with pp1609 linearized Bsm1 (partial digest) to target the plasmid to the *HIS3* locus. Single copy transformants were isolated as described above to generate GPY1890. The *BIM1* and *GIM4* gene knockouts and *TUB1-828-expressing* cassette were then incorporated into GPY1890, as described above, to generate GPY1932 (*Δbim1*) GPY1996 (*Δgim4*) and GPY1935 (*TUB1-828*).

#### STE12 Δste5 strains

To generate these we first needed to construct a plasmid that carried DNA *P_GAL1_* reading STE12 (*GAL1*pr-*STE12*). To do this, we linearized pp1611 with NotI, co-transformed 5ng of the digested plasmid with 20ng of a PCR product that contained the CEN/ARS sequence into BY4741, and plated the transformations on plates deficient in histidine. We prepped 8 isolates using the Zymoprep Yeast Plasmid Miniprep I kit, then transformed into Z-competent *E.coli* to amplify the plasmid, grew single colonies overnight in LB-Amp media and mini-prepped using Qiagen QIAprep Spin Miniprep Kit. Purified plasmids were digested with ApalI to confirm that the CEN/ARS sequence was inserted. We named one such plasmid pSZ99. pSZ99 was then cut with a double digest of ApaI and NcoI to remove the *STE5*-CTM sequence. Next, 5ng of digested pSZ99 and 20ng of *STE12* PCR product (generated using primers GP1110 and GP1111, on BY4741) were co-transformed into BY4741, and transformation mixes plated on histidine deficient media. We isolated DNA from 8 isolates using a Zymoprep Yeast Plasmid Miniprep I kit, then transformed the mix into Z-competent *E.coli*, grew His+ Amp^r^ colonies overnight in LB-Amp media and mini-prepped using Qiagen QIAprep Spin Miniprep Kit. The purified plasmids were then cut with XbaI to verify loss of *STE5*-CTM and insertion of *STE12*, positive isolates were named pSZ98. pSZ98 was then cut with AatII to remove the CEN/ARS, ligated with Rapid T4 DNA Ligase Kit from Roche Diagnostics, transformed into Z-competent *E.col*, and plated on LB-Amp media. 8 colonies were then grown overnight in LB-Amp media and miniprepped using Qiagen QIAprep Spin Miniprep Kit. Purified plasmids were confirmed for CEN/ARS removal by ApaLI digestion, the resulting plasmid was named pSZ97.

GPY1874 was then transformed with pSZ97 linearized with NsiI to target the plasmid to the *HIS3* locus. Single copy transformants were isolated as described above to generate GPY1958. The *BIM1* and *GIM4* gene knockouts and *TUB1-828* were then incorporated into GPY1890, as described above, to generate GPY1976 *(Δbim1)*, GPY1998 (*Δgim4*) and GPY1978 (*TUB1-828*).

### 2.7 *Construction of strains to quantify Ste5 membrane localization*

YPP3662 (Ventura et al.), which carries instead of Ste5 three copies of a Ste5-YFP-YFP-YFP fusion gene, was the the parental strain for these experiments. From it, we generated *Δbim1, Δgim4*, and *tub1-828-expressing* derivatives. We generated GPY4112 (*Δbim1*) and GPY4113 (*Δgim4*), via the same Pringle cassette approach using the URA3(MX4 plasmid) we used to make the GPY4000 series strains. We generated the *tub1-828-expressing* YGP3858 by the same steps we used to generate the strains used in the bypass experiments: transformation by linearized pSZ111 followed by isolation a *P_BMH2_-Gal4-hER-VP16* construct integrated at *BMH2.* We introduced into 8 transformed strains those strains a *P_Gal1_-mCherry* construct, and picked one strain from the majority of strains that showed low expression consistent with single copy insertion of the *P_BMH2_-Gal4-hER-VP16.* We called this strain ACL-GP-001. We then introduced into it pRB2949 (Anders and Botstein, 2001) which carries a P*_GAL1_-TUB1-828* construct, to create ACL-GP-003.

## 3 *Pheromone dose-response experiments*

For follow up studies on the gene deletions of interest and/or all other perturbation experiments, we adapted the high throughput protocol flow cytometry we used for the screen. The only modification was in how we cultured the cells. Instead of culturing them in deep-well 96-well plates, we used the test tube protocol described above for the fluorescence microscopy secondary screen. We did the pheromone stimulation step exactly as in the assays we used in the screen, in shallow-well 96-well plates, with the only difference being the addition of pheromone, which in this case was done using a pipette from a serial dilution of pheromone stocks instead of the slotted-pin tool we used for the screen. All dose response experiments were measured by flow cytometry as described above.

## 4 *Chemical disruption of microtubule polymerization*

We used media with high concentrations of nocodazole (10 μg/ml) and benomyl (30 μg/ml), two chemically distinct but functionally equivalent inhibitors of microtubule polymerization. To prepare the media with these drugs we first heated to 100°C media containing all ingredients for SDC except glucose and buffered with low-strength PBS (0.15 times the standard concentration). We then added 30 μg/ml benomyl from a 10 mg/ml stock in DMSO made daily. We added the same volume of DMSO to a control medium. We then allowed both media to cool to 50 °C, at which point we added 20 g/L glucose and 40 μg/ml casein. Finally we added 10 μg/ml nocodazole from a 10 mg/ml DMSO stock stored in small aliquots at −20 °C. We added the same volume of DMSO to the control media. When indicated, 1-NMPP1 was added to both media to a final concentration of 10 μm.

We tested every batch of nocodazole/benomyl media for its effectiveness at inducing cell cycle arrest in a reference strain. To do so we grew a culture of BY4741 to log phase, spun the cells and resuspended the pellet in the nocodazole/benomyl media. After a 3 hour incubation we counted the number of cells arrested in metaphase, judged by their large budded appearance. The media was considered effective if the fraction of arrested cells was equal or larger than 0.95.

## 5 *5. Consideration of morphology of* Δbim1 *and* Δgim4 *strains*

As we initiated close study in this work of increases in transmitted signal caused by *Δbim1* and *Δgim4* deletions, but before we had carried out the other genetic and chemical perturbation experiments that pointed toward effects on membrane localized, Fus3-dependent signaling, we wished to get a sense of whether the effects of these mutations were on processes at least somewhat specific to signal generation, rather than more general effects due for example to overall disorganization of cell cytoskeletal organization. To this end, we verified by light microscopic observation (not shown) that, compared with reference cells, cells with these lesions showed no great morphological differences during normal growth or after treatment with stimulating pheromone concentrations. Later in this study, we performed, the experiments in Figure 8 to quantify subtle differences in Ste5 signaling patch formation and shmoo formation in *Δbim1* and *Δgim4* cells. In performing those, we again verified by eye that those strains showed no other morphological differences that the ones quantified. Finally, at the suggestion of a reviewer, we subsequently confirmed from published work (Ohya et al., 2005, http://yeast.gLk.u-tokyo.ac.ip)) that *Δbim1* and *Δgim4* cells from the original deletion collection did not show morphological differences from the reference B4741 strain.

## Supplementary Table S1 legend

Strains used in this work.

## References

Armstrong, E. H. (1913) US Patent 1113149, Wireless receiving system, published 19 October 1913, issued 6 October 1914.

Avery, L., and Wasserman, S. (1992). Ordering gene function: the interpretation of epistasis in regulatory hierarchies. Trends Genet 8, 312–316.

Beadle, G. W. and Ephrussi, B. The differentiation of eye pigments in Drosophila as studied by transplantation. Genetics 1936 21, 225–247

Bennett, S. (1979). A history of control engineering. IEEE, Peter Peregrinus, Stevenage, U.K.

Bi, E., and Park, H.O. (2012). Cell polarization and cytokinesis in budding yeast. Genetics 191, 347–387.

Bienz, M., and Clevers, H. (2000). Linking colorectal cancer to Wnt signaling. Cell 103, 311–320.

Botstein, D., and Maurer, R. (1982). Genetic approaches to the analysis of microbial development. Annual Rev. Genet. 16, 61–83.

Botstein, D., D. Amberg, D. Mulholland, J. Huffaker, T., Adams, A. Drubin, D and Stearns, T. The Yeast Cytoskeleton (1997). In the molecular and cellular biology of the yeast Saccharomyces. J.R. Broach, J.R. Pringle, and E.W. Jones, editors. 1997. Cold Spring Harbor Laboratory Press, Cold Spring Harbor, NY. 1–90.

Brenner, S. (2010). Sequences and consequences. Phil. Trans. R. Soc B. (2010) 365, 207–212

Brent, R. (2009). Cell signaling: what is the signal and what information does it carry? FEBS Lett. 583, 4019–4024.

Brent, R., and Finley, R.L.,Jr. (1997). Understanding gene and allele function with two-hybrid methods. Ann. Rev. Genet. 31, 663–704.

Bush, A., Chernomoretz, A., Yu, R., Gordon, A., and Colman-Lerner, A. (2012). Using Cell-ID 1.4 with R for microscope-based cytometry. Current protocols in molecular biology / edited by Frederick M Ausubel, Roger Brent, et al. Chapter 14, Unit 14 18.

Bush, A., and Colman-Lerner, A. (2013). Quantitative Measurement of Protein Relocalization in Live Cells. Biophys. J. 104(3): 727–736

Butty, A.C., Perrinjaquet, N., Petit, A., Jaquenoud, M., Segall, J.E., Hofmann, K., Zwahlen, C., and Peter, M. (2002). A positive feedback loop stabilizes the guanine-nucleotide exchange factor Cdc24 at sites of polarization. EMBO J. 21, 1565–1576.

Butty, A.C., Pryciak, P.M., Huang, L.S., Herskowitz, I., and Peter, M. (1998). The role of Far1p in linking the heterotrimeric G protein to polarity establishment proteins during yeast mating. Science 282, 1511–1516.

Cairns B. R., Ramer, S. W., and Kornberg, R. D. (1992). Order of action of components in the yeast pheromone response pathway revealed with a dominant allele of the STE11 kinase and the multiple phosphorylation of the STE7 kinase. Genes Dev. 6(7): 1305–1318.

Casolari, J.M., Brown, C.R., Drubin, D.A., Rando, O.J., and Silver, P.A. (2005). Developmentally induced changes in transcriptional program alter spatial organization across chromosomes. Genes Dev 19, 1188–1198.

Cheong, R., Rhee, A., Wang, C.J., Nemenman, I., and Levchenko, A. (2011). Information transduction capacity of noisy biochemical signaling networks. Science 334, 354–358.

Colman-Lerner, A., Gordon, A., Serra, E., Chin, T., Resnekov, O., Endy, D., Pesce, C.G., and Brent, R. (2005). Regulated cell-to-cell variation in a cell-fate decision system. Nature 437, 699–706.

Conlon, P., Gelin-Licht, R., Ganesan, A., Zhang, J., and Levchenko, A. (2016). Single-cell dynamics and variability of MAPK activity in a yeast differentiation pathway. Proc. Natl. Acad. Sci. USA. 113(40): E5896–E5905.

Dolan, J.W., and Fields, S. (1990). Overproduction of the yeast STE12 protein leads to constitutive transcriptional induction. Genes Dev. 4, 492–502.

Drogen, F., O’Rourke, S.M., Stucke, V.M., Jaquenoud, M., Neiman, A.M., and Peter, M. (2000). Phosphorylation of the MEKK Ste11p by the PAK-like kinase Ste20p is required for MAP kinase signaling in vivo. Curr. Biol. 10, 630–639.

Efron, B. (1979). Bootstrap methods: another look at the jackknife. Annals Statistics, 7, 1–26.

Erlemann, S., Neuner, A., Gombos, L., Gibeaux, R., Antony, C, and Schiebel, E. (2012). An extended γ-tubulin ring functions as a stable platform in microtubule nucleation. J. Cell. Biol. 197, 29–74

Evangelista, M., Blundell, K., Longtine, M.S., Chow, C.J., Adames, N., Pringle, J.R., Peter, M., and Boone, C. (1997). Bni1p, a yeast formin linking cdc42p and the actin cytoskeleton during polarized morphogenesis. Science 276, 118–122.

Fu, W., O’Connor, T.D., Jun, G., Kang, H.M., Abecasis, G., Leal, S.M., Gabriel, S., Rieder, M.J., Altshuler, D., Shendure, J., et al. (2013). Analysis of 6,515 exomes reveals the recent origin of most human protein-coding variants. Nature 493, 216–220.

Gärtner, K. (1990). A third component causing random variability beside environment and genotype. A reason for the limited success of a 30 year long effort to standardize laboratory animals? Lab. Anim. 24: 71–77.

Geissler, S., Siegers, K., and Schiebel, E. (1998). A novel protein complex promoting formation of functional alpha- and gamma-tubulin. EMBO J. 17, 952–966.

Gelin-Licht, R., Paliwal, S., Conlon, P., Levchenko, A., and Gerst, J. E. (2012). Scp160-dependent mRNA trafficking mediates pheromone gradient sensing and chemotropism in yeast. Cell Rep. 1: 483–494.

Good, M., Tang, G., Singleton, J., Remenyi, A., and Lim, W.A. (2009). The Ste5 scaffold directs mating signaling by catalytically unlocking the Fus3 MAP kinase for activation. Cell 136, 1085–1097.

Good, P.I. (2004). Permutation, Parametric, and Bootstrap Tests of Hypotheses (Springer Series in Statistics) (Springer-Verlag New York, Inc.).

Gordon, A., Colman-Lerner, A., Chin, T.E., Benjamin, K.R., Yu, R.C., and Brent, R. (2007). Single-cell quantification of molecules and rates using open-source microscope-based cytometry. Nat Methods 4, 175–181.

Gundersen, G.G., and Worman, H.J. (2013). Nuclear positioning. Cell 152, 1376–1389.

Harris, K., Lamson, R.E., Nelson, B., Hughes, T.R., Marton, M.J., Roberts, C.J., Boone, C., and Pryciak, P.M. (2001). Role of scaffolds in MAP kinase pathway specificity revealed by custom design of pathway-dedicated signaling proteins. Curr. Biol, 11, 1815–1824.

Hodgkin, J. (2005) Genetic suppression, WormBook, ed. The *C. elegans* Research Community, WormBook, December 7, 2005, doi/10.1895/wormbook.1.59.1, http://www.wormbook.org

Irazoqui, J. E., Gladfelter, A. S., and Lew, D. J. (2003) Scaffold-mediated symmetry breaking by Cdc42p. Nat. Cell Biol. 5(12): 1062–1070

Jackson, C.L., and Hartwell, L.H. (1990). Courtship in *S. cerevisiae:* both cell types choose mating partners by responding to the strongest pheromone signal. Cell 63, 1039–1051.

Jimenez-Gomez, J. M., Corwin, J. A.; Joseph, B., Maloof, J. N., Kliebenstein, D. J. (2011) Genomic analysis of QTLs and genes altering natural variation in stochastic noise. PLOS Genet. 7 (9): e1002295. 5.

Johnson, J.M., Jin, M., and Lew, D.J. (2011). Symmetry breaking and the establishment of cell polarity in budding yeast. Curr. Opin. Genet. Dev. 21, 740–746.

Karpova, T.S., Reck-Peterson, S. L., Elkind, N. B., Mooseker, M. S., Novick, P. J., and Cooper, J.A. (2000). Role of actin and Myo2p in polarized secretion and growth of *Saccharomyces cerevisiae.* Mol. Biol. Cell 11, 1727–1737.

Kinzler, K.W., and Vogelstein, B. (1996). Lessons from hereditary colorectal cancer. Cell 87, 159–170.

Kozubowski, L., Saito, K., Johnson, J. M., Howell, A. S., Zyla, T. R., and Lew, D. J. (2008) Symmetry-breaking polarization driven by a Cdc42p GEF-PAK complex. Curr Biol. 18(22): 1719–1726.

Lesko, A.C., Goss, K.H., and Prosperi, J.R. (2014). Exploiting APC function as a novel cancer therapy. Current Drug Targets 15, 90–102.

Lipkow, K., Andrews, S.S., and Bray, D. (2005). Simulated diffusion of phosphorylated CheY through the cytoplasm of Escherichia coli. J. Bacteriol. 187, 45–53.

Louvion, J. F., Havaux-Copf, B., and Picard, D. (1993). Fusion of GAL4-VP16 to a steroid-binding domain provides a tool for gratuitous induction of galactose-responsive genes in yeast. Gene 131(1): 129–134

Maddox, P., Chin, E., Mallavarapu, A., Yeh, E., Salmon, E. D., and Bloom K. (1999). Microtubule dynamics from mating through the first zygotic division in the budding yeast Saccharomyces cerevisiae. J Cell Biol. 144, 977–987.

Maddox, P.S., Stemple, J.K., Satterwhite, L., Salmon, E.D., and Bloom, K. (2003). The minus end-directed motor Kar3 is required for coupling dynamic microtubule plus-ends to the cortical shmoo tip in budding yeast. Curr. Biol. 13, 1423–1428.

Maeder, C.I., Hink, M.A., Kinkhabwala, A., Mayr, R., Bastiaens, P.I., and Knop, M. (2007). Spatial regulation of Fus3 MAP kinase activity through a reaction-diffusion mechanism in yeast pheromone signalling. Nat. Cell Biol. 9, 1319–1326.

Makumburage, G. B., and Stapleton, Ann E. (2011). Phenotype Uniformity in Combined-Stress Environments has a Different Genetic Architecture than in Single-Stress Treatments. In Frontiers in Plant Science 2: 12

Meiksin, Z. H. (1959). Positive-feedback phase-space trajectories and application to servo systems. IEEE Electrical Engineering 78, 673–679

Meiksin, Z. H. (1959). Positive feedback in servomechanisms. IEEE Electrical Engineering, 78, 1095

Meissner, D. E. (1913) German Patent 291604, Einrichtung zur Erzeugung elektrischer Schwingungen, issued to Gesellschaft für Drahtlose Telegraphie mbH, published April 10, 1913, issued June 23, 1919

Molk, J. N., and Bloom, K. (2006) Microtubule dynamics in the budding yeast mating pathway. J. Cell Sci. 119(Pt 17): 3485–3490.

Molk, J. N., Salmon, E. D., and Bloom, K. (2006) Nuclear congression is driven by cytoplasmic microtubule plus end interactions in *S. cerevisiae*. J. Cell Biol. 172(1): 27–39.

Moskow J. J., Gladfelter, A. S., Lamson, R. E., Pryciak, P. M., and Lew, D. J. (2000) Role of Cdc42p in pheromone-stimulated signal transduction in *Saccharomyces cerevisiae*. Mol. Cell Biol. 20(20): 7559–7571.

Murphy, H. A. and Zeyl, C. W. (2010) Yeast sex: surprisingly high rates of outcrossing between asci. PLoS ONE. 5: e10461.

Murphy, H. A., and Zeyl, C. W. (2012). Prezygotic isolation between *Saccharomyces cerevisiae* and *Saccharomyces paradoxus* through differences in mating speed and germination timing. Evolution. 66: 1196–1200

Murphy, H. A., H. A. Kuehne, C. A. Francis, and P. D. Sniegowski (2006) Mate choice assays and mating propensity differences in natural yeast populations. Biol. Lett. 2: 553–556.

Nern, A., and Arkowitz, R.A. (1999). A Cdc24p-Far1p-Gbetagamma protein complex required for yeast orientation during mating. J. Cell Biol. 144, 1187–1202.

Novick P., Ferro S. and Schekman, R. (1981) Order of events in the yeast secretory pathway. Cell 25: 461–469.

Palframan, W. J., Meehl, J. B., Jaspersen, S. L., Winey, M., and Murray, A. W (2006). Anaphase inactivation of the spindle checkpoint. Science 313: 680–684.

Paterson, J. M., Ydenberg, C. A., and Rose, M. D. (2008). Dynamic localization of yeast Fus2p to an expanding ring at the cell fusion junction during mating. J. Cell Biol. 181(4): 697–709.

Pruyne, D. (2001). Finding direction: the ins and outs of cell polarity. J. Cell Sci. 114, 5–8.

Pryciak, P.M., and Huntress, F.A. (1998). Membrane recruitment of the kinase cascade scaffold protein Ste5 by the Gβγ complex underlies activation of the yeast pheromone response pathway. Genes Dev. 12, 2684–2697.

Read, E.B., Okamura, H.H., and Drubin, D.G. (1992). Actin- and tubulin-dependent functions during Saccharomyces cerevisiae mating projection formation. Mol. Biol. Cell 3, 429–444.

Sayas, C.L., and Avila, J. (2014). Regulation of EB1/3 proteins by classical MAPs in neurons. Bioarchitecture 4, 1–5.

Schmalhausen, I. I. (1949): Factors of evolution: the theory of stabilizing selection. University of Chicago Press, Chicago, reprint of publication by Blakiston Co., Philedelphia

Song, D., Dolan, J.W., Yuan, Y.L., and Fields, S. (1991). Pheromone-dependent phosphorylation of the yeast STE12 protein correlates with transcriptional activation. Genes Dev. 5, 741–750.

Spencer, S.L., Gaudet, S., Albeck, J.G., Burke, J.M., and Sorger, P.K. (2009). Non-genetic origins of cell-to-cell variability in TRAIL-induced apoptosis. Nature 459, 428–432.

Stone, E.M., Heun, P., Laroche, T., Pillus, L., and Gasser, S.M. (2000). MAP kinase signaling induces nuclear reorganization in budding yeast. Curr. Biol. 10, 373-382.

Stransky, N., Egloff, A. M., Tward, A. D., Kostic, A. D., Cibulskis, K., Sivachenko, A., Kryukov, G. V., Lawrence, M. S., Sougnez, C., McKenna, A., et al. (2011). The mutational landscape of head and neck squamous cell carcinoma. Science 333, 1157–1160.

Su, L. K., and Qi, Y. (2001). Characterization of human MAPRE genes and their proteins. Genomics 71, 142–149.

Suchkov, D.V., DeFlorio, R., Draper, E., Ismael, A., Sukumar, M., Arkowitz, R., and Stone, D.E. (2010). Polarization of the yeast pheromone receptor requires its internalization but not actindependent secretion. Mol. Biol. Cell 21, 1737–1752.

Takahashi, S., and Pryciak, P.M. (2008). Membrane localization of scaffold proteins promotes graded signaling in the yeast MAP kinase cascade. Curr. Biol. 18, 1184–1191.

Tedford, K., Kim, S., Sa, D., Stevens, K., and Tyers, M. (1997). Regulation of the mating pheromone and invasive growth responses in yeast by two MAP kinase substrates. Curr. Biol. 7, 228–238.

Ten Hoopen, R., Cepeda-García, C., Fernández-Arruti, R., Juanes, M. A., Delgehyr, N., and Segal M. (2012). Mechanism for astral microtubule capture by cortical Bud6p priming spindle polarity in *S. cerevisiae*. Curr. Biol. 22(12): 1075–1083.

Thomson, T.M., Benjamin, K.R., Bush, A., Love, T., Pincus, D., Resnekov, O., Yu, R.C., Gordon, A., Colman-Lerner, A., Endy, D., and Brent, R. (2011). Scaffold number in yeast signaling system sets tradeoff between system output and dynamic range. Proc. Natl. Acad. Sci. USA 108, 20265–20270.

Valtz, N., Peter, M., and Herskowitz, I. (1995). FAR1 is required for oriented polarization of yeast cells in response to mating pheromones. J. Cell. Biol. 131, 863–873.

van Drogen, F., Stucke, V.M., Jorritsma, G., and Peter, M. (2001). MAP kinase dynamics in response to pheromones in budding yeast. Nat. Cell. Biol. 3, 1051–1059.

Ventura, A. C., Bush, A., Vasen, G., Goldín, M. A., Burkinshaw, B., Bhattacharjee, N., Folch, A., Brent, R., Chernomoretz, A., and Colman-Lerner A. (2014) Utilization of extracellular information before ligand-receptor binding reaches equilibrium and expands and shifts the input dynamic range. Proc. Natl. Acad. Sci. USA. 111(37): e3860–3869

Vogelstein, B., Papadopoulos, N., Velculescu, V.E., Zhou, S., Diaz, L.A., Jr., and Kinzler, K.W. (2013). Cancer genome landscapes. Science 339, 1546–1558.

Wuestehube L. J., Duden, R., Eun, A., Hamamoto, S., Korn, P., Ram, R., and Schekman, R. (1996). New mutants of Saccharomyces cerevisiae affected in the transport of proteins from the endoplasmic reticulum to the Golgi complex. Genetics: 142(2): 393–406

Yi, T.M., Kitano, H., and Simon, M.I. (2003). A quantitative characterization of the yeast heterotrimeric G protein cycle. Proc Natl Acad Sci USA 100, 10764–10769.

Yokota, Y., Kim, W.Y., Chen, Y., Wang, X., Stanco, A., Komuro, Y., Snider, W., and Anton, E.S. (2009). The adenomatous polyposis coli protein is an essential regulator of radial glial polarity and construction of the cerebral cortex. Neuron 61, 42–56.

Yu, R.C., Pesce, C.G., Colman-Lerner, A., Lok, L., Pincus, D., Serra, E., Holl, M., Benjamin, K., Gordon, A., and Brent, R. (2008). Negative feedback that improves information transmission in yeast signalling. Nature 456, 755–761.

Zaichick, S. V., Metodiev, M. V., Nelson, S. A., Durbrovskyi, O., Draper, E., Cooper, J. A., and Stone DE. (2009). The mating-specific Gα interacts with a kinesin-14 and regulates pheromone-induced nuclear migration in budding yeast. Mol Biol Cell 20: 2820–2830.

## References

Huh, W.-K., Falvo, J. V., Gerke, L. C., Carroll, A. S., Howson, R. W., Weissman, J. S., an O'Shea, E.K (2003). Global analysis of protein localization in budding yeast. Nature. 425(6959): 686–691

Longtine, M. S., McKenzie, A. 3rd, Demarini, D. J., Shah, N. G., Wach, A., Brachat, A., Philippsen, P., and Pringle J.R. (1998). Additional modules for versatile and economical PCR-based gene deletion and modification in *Saccharomyces cerevisiae*. Yeast 14(10): 953–961.

Louvion, J. F., Havaux-Copf, B., and Picard, D. (1993) Fusion of GAL4-VP16 to a steroid-binding domain provides a tool for gratuitous induction of galactose-responsive genes in yeast. Gene. 131: 129–134

Ohya, Y., Sese, J., Yukawa, M., Sano, F., Nakatani, Y., Saito, T. L., Saka, A., Fukuda, T., Ishihara, S., Oka, S., Suzuki, G., Watanabe, M., Hirata, A., Ohtani, M., Sawai, H., Fraysse, N., Latgé, J. P., François, J. M., Aebi, M., Tanaka, S., Muramatsu, S., Araki, H., Sonoike, K., Nogami, S., and Morishita, S. (2005). High-dimensional and large-scale phenotyping of yeast mutants. Proc. Natl. Acad. Sci. USA. 102(52): 19015–19020.

Pesce, C., G., Zdraljevic, S., 3, Peria, W., Rockwell, D., Yu, R. C., Colman-Lerner, A., and Brent, R. (2016). Cell-to-cell variability in the yeast pheromone response: high throughput screen identifies genes with different effects on transmitted signal and response. Molecular and Systems Biology, submitted

Straight, A. F., Sedat, J. W., and Murray, A. W. (1998) Time-lapse microscopy reveals unique roles for kinesins during anaphase in budding yeast. J. Cell Biol. 143: 687–694

Ventura, A. C., Bush, A., Vasen, G., Goldín, M. A., Burkinshaw, B., Bhattacharjee, N., Folch, A., Brent, R., Chernomoretz, A., and Colman-Lerner, A. (2014) Utilization of extracellular information before ligand-receptor binding reaches equilibrium expands and shifts the input dynamic range. Proc. Natl. Acad. Sci. USA. 111(37): e3860–e3869

